# Pseudocryptic diversity within the *Phakellia robusta* species complex: description of *Phakellia pallida* sp. nov. and *Phakellia pusilla* sp. nov

**DOI:** 10.64898/2026.10.06.756958

**Authors:** Carlota Gracia-Sancha, Sergi Taboada, Ana De la Torriente, Carles Galià-Camps, Marcos González-Porto, Pilar Ríos, Javier Cristobo, Alberto Serrano, Enric Massutí, Ana Riesgo, Julio A. Díaz

## Abstract

The genus *Phakellia* comprises predominantly fan-shaped deep-sea sponges, but its species diversity and boundaries remain unresolved. Here, we use an integrative approach combining morphological and molecular data to investigate the *Phakellia robusta* species complex and describe two new species, ***Phakellia pallida* sp. nov**. and ***Phakellia pusilla* sp. nov**. Overlap in external and spicular morphology makes these taxa challenging to identify. Molecular analyses based on mitochondrial (*cox1*) and nuclear (*28S*) markers support the recognition of the three taxa as distinct species, and reveal their close evolutionary relationship. The two new species co-occur in the NE Atlantic and the Mediterranean, with overlapping depth ranges, although *P. pallida* sp. nov. is generally found at greater depths. Several previous records of *P. robusta* from the Mediterranean and southern NE Atlantic may instead correspond to one of the two newly described species, indicating that its distribution requires reassessment. A group I intron previously reported in *P. robusta* was also detected at the same *cox1* insertion site in some, but not all, individuals of *P. pusilla* sp. nov., but was absent from *P. pallida* sp. nov. Overall, our results reveal pseudocryptic diversity within *Phakellia* and highlight the value of integrative approaches for resolving species boundaries.

## Introduction

Species delimitation has traditionally relied on morphologically-based classification systems. However, speciation is not always accompanied by clear morphological differentiation, and genetically distinct lineages may remain hidden under a single nominal species (Avise et al., 1994). In marine environments, identifying biodiversity is particularly challenging because genetically distinct lineages often exhibit little or no morphological differentiation, due to morphological stasis or convergent evolution driven by similar ecological pressures, resulting in cryptic diversity (Bickford *et al*., 2007; Hending, 2025). In some cases, however, genetically distinct lineages initially regarded as cryptic can subsequently be distinguished by subtle but consistent morphological characters, revealing pseudocryptic rather than truly cryptic diversity (Knowlton 1993; Bickford *et al*., 2007). Moreover, reproductive isolation in many marine organisms may rely more on chemical, temporal, or ecological signals than on visually recognizable traits, potentially reducing the selective pressures for conspicuous morphological divergence (Knowlton, 2000; Bickford *et al*., 2007). Consequently, morphologically similar organisms may hide substantial genetic divergence, obscuring species boundaries and leading to an underestimation of marine biodiversity.

Sponges (Porifera) represent one of the best-known examples of hidden marine diversity. Currently, there are around 9,875 sponge species described (de Voogd *et al*., 2026), with estimates of 15,000 species in total (van Soest *et al*., 2012). The lack of diagnostic external characters, combined with widespread morphological convergence and a long tradition of morphology-based taxonomy, has obscured species boundaries and masked substantial genetic diversity (Hooper & Van Soest, 2002). Morphological characters, particularly the size and shape of spicules, can sometimes help to distinguish closely related species (Cárdenas *et al*., 2007). However, there are also cases where neither external nor internal morphology reflects underlying genetic variation (Klautau *et al.,* 1999). Therefore, the application of molecular tools, including DNA markers, allozymes, and microbiome analyses, has revealed that many sponge species previously considered cosmopolitan or widely distributed, actually comprise complexes of cryptic species, particularly in Demospongiae (e.g. Muricy et al., 1996; Xavier et al., 2010; de Paula et al., 2012; DeBiasse & Hellberg, 2015; Uriz et al., 2017; Swierts et al., 2017; Pankey et al., 2024). Among demosponges, the genus *Phakellia* Bowerbank, 1862 represents an interesting case for investigating the potential occurrence of cryptic diversity due to its complex taxonomic history, the paucity of diagnostic characters for the genus, and the increasing availability of molecular data suggesting morphological and genetic discordance (Morrow & Cárdenas, 2015; Taboada *et al*., 2022). *Phakellia* is a genus within the Bubarida order, composed of fan-shaped sponges encompassing 33 recognized species (de Voogd *et al*., 2026), mostly associated with mesophotic and deep sea rocky environments, including banks and canyons (Topsent, 1928; Ríos *et al*., 2017; Sánchez *et al*., 2017; Taboada *et al*., 2022, 2023). The genus exhibits a wide geographic and bathymetric distribution, occurring from shallow to deep waters in tropical, temperate, and polar regions worldwide. It shows particularly diversification in the Mediterranean Sea but is also found throughout the Atlantic, Indian, and Pacific Oceans (van Soest, 1994; Alvarez & Hooper, 2002). Despite the broad ecological range, several species act as keystone components of deep-sea sponge grounds in the North Atlantic (Maldonado *et al*., 2016; Taboada *et al*., 2022).

The genus *Phakellia* has long suffered from taxonomic intricacies due to the lack of clear synapomorphies for the group. Macroscopic morphology is characterised by pedunculate forms ranging from flabellate fan to cup shaped sponges, with orange to white colors (Alvarez *et al*., 2016; Santín *et al*., 2018; Gayá-Vilar *et al*., 2024). Spicule types include sinuous or vermiform diactinal spicules with rounded to sharp ends, which are called oxeas and styles (adapted from Alvarez & Hooper, 2002), a spicule complement that is shared between distant sponge clades, such as vilesids and axinellids (Díaz *et al.,* 2025). This paucity of characters has fueled their complex taxonomic history. Traditionally, Phakellia was placed in the order Axinellida and family Axinellidae Carter, 1875 based on their similar morphological and spicular features (Alvarez & Hooper, 2002). However, molecular phylogenetic studies demonstrated that Axinellidae was a polyphyletic group (Morrow and Cárdenas 2015). As a result of the taxonomic revision proposed by Morrow and Cárdenas (2015), the genus was transferred to the family Bubaridae Topsent, 1894 within the new order Bubarida (Morrow & Cárdenas, 2015), where it currently remains. Therefore, there has been a reclassification of many species previously described in the genus *Phakellia* later reassigned to other genera, being *Axinella* Schmidt, 1862 and *Acanthella* (Schmidt, 1862) the more common new placements (Hooper & Levi, 1993).

Despite the revisions in the family Axinellidae, important taxonomic uncertainties remain unresolved. A notable example is the case of *Phakellia robusta* Bowerbank, 1866, a species with a broad depth range (20-1,229 m) and a distribution spanning the NE Atlantic Ocean and the Mediterranean Sea (Topsent, 1904, 1925, 1928; Ferrer Hernández, 1914; Stephens, 1921; Alander, 1942; Vacelet, 1969; Maldonado, 1992; Cruz, 2002; Ríos *et al*., 2017; 2022; Santín *et al*., 2018). Morphologically, this species is described as a fan-shaped sponge attached by a short peduncle. Minor morphological variations have been reported both externally, with specimens showing the presence or absence of longitudinal striations extending from the base to the lamellar margin, and internally, with vermiform spicules exhibiting either rounded or pointed ends depending on the individual (Bowerbank, 1866; Topsent, 1904, 1928; Taboada *et al*., 2022). Genetically, phylogenetic analyses have positioned individuals from the Mediterranean and the Cantabrian Sea as distinct from those of the NE Atlantic ones, supporting the designation of a new taxonomic entity, *Phakellia* aff. *robusta* (Díaz *et al*., 2025). Moreover, the discovery of hybrids between *Phakellia robusta* and *Phakellia* aff. *robusta* (Taboada *et al*., 2022) (formerly *Phakellia hirondellei*) further challenges traditional species boundaries. Taken together, these morphological inconsistencies, the potential hybrids, and the slightly different distribution of distinct genetic lineages suggest the existence of a species complex within *P. robusta* and highlight the need for a comprehensive systematic revision of the group (Díaz *et al*., 2025).

Resolving taxonomic inconsistencies in this genus is important, as some *Phakellia* species (including *P. robusta*) are known to form sponge grounds in mesophotic and deep sea habitats around the NE Atlantic Ocean and the Mediterranean (Kutti *et al*., 2013; Maldonado *et al*., 2016; Sánchez *et al*., 2017; Díaz *et al*., 2024; Frank *et al.,* 2026). These ecosystems are key components of deep sea habitats, being recognized as hotspots of biodiversity, providing habitat and shelter for numerous marine organisms while enhancing local biodiversity, driving nutrient cycling and promoting benthic-pelagic coupling (De Goeij *et al*., 2013; Cathalot *et al*., 2015; Maldonado *et al*., 2016; Rix *et al*., 2018). Owing to their key role in structuring benthic communities and shaping the seafloor, these species have become the focus of increasing ecological, genetic, and genomic research, improving our understanding of their diversity, evolutionary relationships, population connectivity, and contribution to deep-sea ecosystem functioning (Cranston *et al*., 2021; Taboada *et al*., 2022, 2023; Rodríguez-Basalo *et al*., 2022; Koutsouveli *et al*., 2022). In this scenario, taxonomic uncertainties within *Phakellia* remain a major challenge for their effective conservation and management. Misidentification, together with the likely occurrence of cryptic species within the genus, can lead to inaccurate biodiversity assessments and inappropriate management decisions (Bickford *et al*., 2007; Robinson *et al*., 2014). As morphology-based species identification remains particularly challenging in *Phakellia*, the present study aims to resolve the *P. robusta* species complex using an integrative taxonomic approach that combines morphological and molecular data, thereby clarifying species boundaries within the complex and reassessing the phylogenetic relationships of the genus *Phakellia*.

## Material and methods

### Sampling

Material collected (Supplementary Table 1) originates from research surveys conducted in the North Atlantic Ocean, the Balearic Islands (western Mediterranean) and the Cantabrian Sea (NE Atlantic). These surveys were carried out during the IEO-CSIC campaigns LIFE IP INTEMARES between 2019 and 2020 (Massutí *et al*., 2022), the MEDITS surveys between 2016 and 2021, CIRCAROCK and CIRCALEBA cruises, conducted between May and June from 2022 to 2024 under the Spanish Marine Strategy Framework Directive (MSFD - project ESMARES-CIRCA) monitoring program (Long, 2011; Bellas, 2014), and the SponGES project in June 2017. Other Atlantic samples were collected from the Canary Islands during the IEO-CSIC CANZEC cruise in 2021, from Hatton Rockall Basin on board the MFV Scotia, July 2015, and from the Røst Reef during the Polarstern Expedition ARK XII/1a in 2007. Beam trawls (BT), the experimental GOC-73 trawl, rock dredges (RD), anchor dredge (AD) or remotely operated vehicles (ROV) were used for the sponge collection. Whole specimen voucher samples were preserved on deck in 96% EtOH. In parallel, small pieces (1 cm^3^) of each specimen were cut and preserved in 96% EtOH for molecular analyses, on which the ethanol was replaced after 24 hours to ensure a better preservation of the DNA. Documentation and photography of sponge specimens were conducted *in situ* (ROV) and on-deck whenever possible. Type specimens of *P. robusta* Bowerbank, 1866 from the Natural History Museum zoological collection and comparative material from *P. ventilabrum* from the Cantabrian and the Mediterranean Sea were also revised (Fig. 1; Table 1).

**Figure 1.**
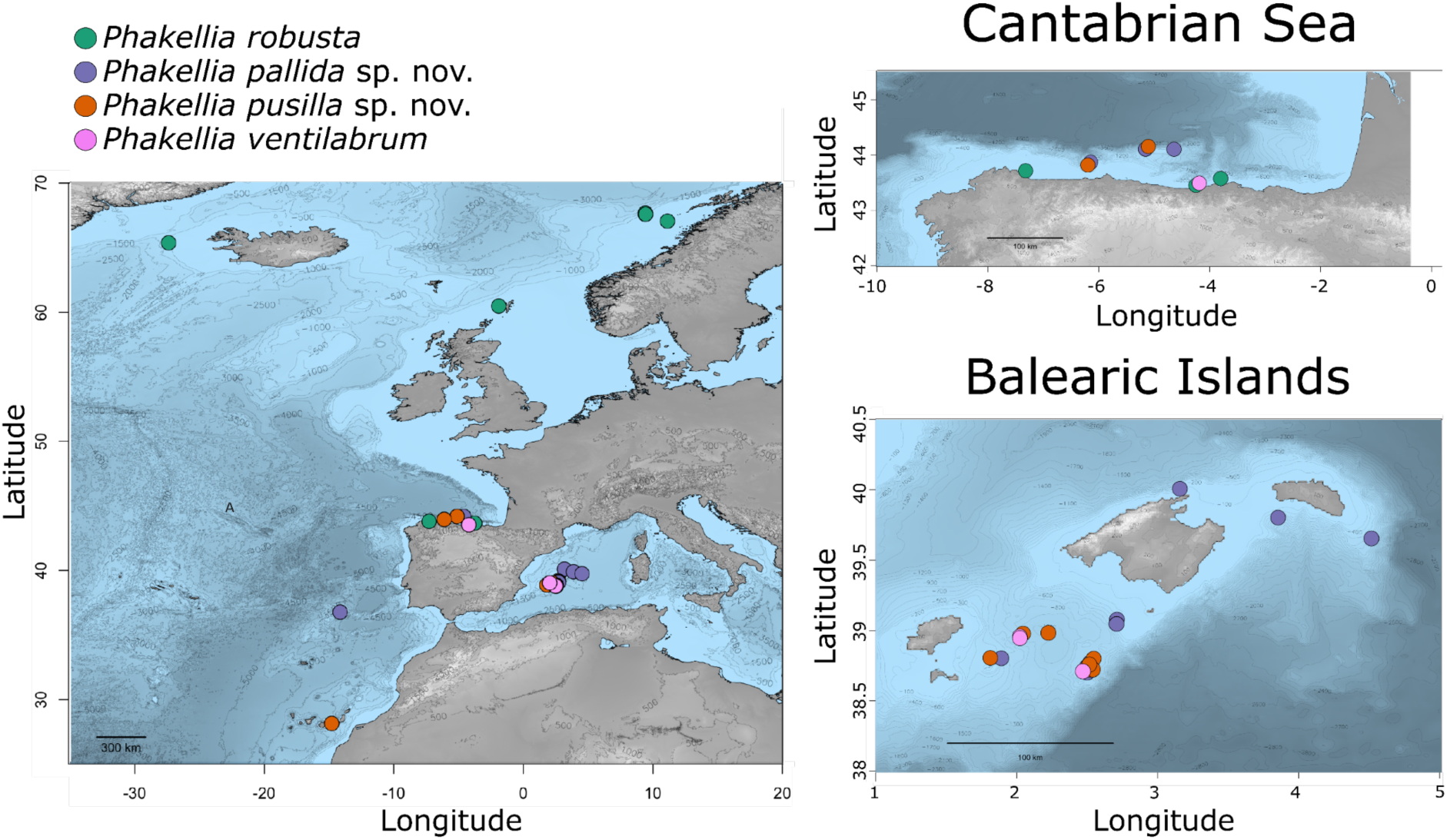
Map indicating the sampling locations where specimens of the genus *Phakellia* were collected; each color represents a different species. Top right: detail of the sampling locations in the Cantabrian Sea (NE Atlantic). Bottom right: detail of the sampling locations in the Balearic Islands (W Mediterranean).

**Table 1.** List of specimens used in the study, organized by species, with sampling information including area, region, depth, latitude and longitude.

| Species/Code | Area | Region | Depth (m) | Latitude | Longitude |
| --- | --- | --- | --- | --- | --- |
| *Phakellia robusta* |  |  |  |  |  |
| 27_1-1 | NE Atlantic | Røst Reef | 292 | 66.9720 | 11.1167 |
| 14_4-14 | NE Atlantic | Røst Reef | 300 | 67.5000 | 9.4033 |
| 14_4-13 | NE Atlantic | Røst Reef | 300 | 67.5000 | 9.4033 |
| 14_4-17 | NE Atlantic | Røst Reef | 300 | 67.6000 | 9.4033 |
| 14_4-19 | NE Atlantic | Røst Reef | 300 | 67.6000 | 9.4033 |
| 9_4-3 | NE Atlantic | Røst Reef | 300 | 67.5083 | 9.4267 |
| 191055 | NE Atlantic | Denmark Strait | 468–565 | 65.2900 | -27.4567 |
| ST6_A | NE Atlantic | Korsfjorden | 29–213 | 59.8728 | 5.5499 |
| ST2_30a | NE Atlantic | Korsfjorden | 29–213 | 59.8728 | 5.5499 |
| KHR13 | NE Atlantic | Kerry Head Reefs | 105 | 52.3430 | -10.7350 |
| M232 | NE Atlantic | Cantabrian Sea | 96 | 43.5742 | -3.7946 |
| M296 | NE Atlantic | Cantabrian Sea | 75 | 43.4515 | -4.2411 |
| M70 | NE Atlantic | Cantabrian Sea | 50 | 43.7135 | -7.3156 |
| 420 | NE Atlantic | Shetland Islands | - | 60.3954 | -1.9276 |
| *Phakellia pallida* sp. nov. |  |  |  |  |  |
| MON33 | NE Atlantic | Josephine Bank | 208 | 36.6833 | -14.1958 |
| ES817 | W Mediterranean | Balearic Islands | 211 | 40.0147 | 3.1505 |
| POR267 | W Mediterranean | Balearic Islands | 248 | 39.0821 | 2.7034 |
| POR462 | W Mediterranean | Balearic Islands | 355 | 39.8080 | 3.8476 |
| POR760 | W Mediterranean | Balearic Islands | 252 | 39.0510 | 2.7008 |
| POR1050 | W Mediterranean | Balearic Islands | 674 | 39.6597 | 4.5108 |
| POR1231 | W Mediterranean | Balearic Islands | 355 | 39.0498 | 2.7007 |
| i288 | W Mediterranean | Balearic Islands | 336–458 | 38.8047 | 1.8817 |
| i347_0 | W Mediterranean | Balearic Islands | 170 | 38.7452 | 2.4905 |
| i347_2 | W Mediterranean | Balearic Islands | 170 | 38.7452 | 2.4905 |
| i409_1 | W Mediterranean | Balearic Islands | 151 | 38.7062 | 2.4937 |
| i414_3 | W Mediterranean | Balearic Islands | 412 | 38.7727 | 2.5125 |
| i713_4 | W Mediterranean | Balearic Islands | 147 | 38.7050 | 2.4908 |
| i713_5 | W Mediterranean | Balearic Islands | 147 | 38.7050 | 2.4908 |
| DR7_253 | NE Atlantic | Cantabrian Sea: El Cachucho | 660 | 44.1034 | -5.1510 |
| DR9_400 | NE Atlantic | Cantabrian Sea: El Cachucho | 660 | 44.0450 | -5.1057 |
| DR10_455 | NE Atlantic | Cantabrian Sea: El Cachucho | 541 | 44.1013 | -4.6383 |
| DR10_456 | NE Atlantic | Cantabrian Sea: El Cachucho | 541 | 44.1013 | -4.6383 |
| DR10_457 | NE Atlantic | Cantabrian Sea: El Cachucho | 541 | 44.1013 | -4.6383 |
| DR10_458 | NE Atlantic | Cantabrian Sea: El Cachucho | 541 | 44.1013 | -4.6383 |
| DR10_459 | NE Atlantic | Cantabrian Sea: El Cachucho | 541 | 44.1013 | -4.6383 |
| DR10_464 | NE Atlantic | Cantabrian Sea: El Cachucho | 541 | 44.1013 | -4.6383 |
| DR10_466 | NE Atlantic | Cantabrian Sea: El Cachucho | 541 | 44.1013 | -4.6383 |
| DR10_498 | NE Atlantic | Cantabrian Sea: El Cachucho | 541 | 44.1013 | -4.6383 |
| DR14_652 | NE Atlantic | Cantabrian Sea: Avilés Canyon System | 340 | 43.8680 | -6.1425 |
| DR15_857b | NE Atlantic | Cantabrian Sea: El Cachucho | 650 | 44.1057 | -5.1505 |
| DR15_857c | NE Atlantic | Cantabrian Sea: El Cachucho | 650 | 44.1057 | -5.1505 |
| DR15_976 | NE Atlantic | Cantabrian Sea: El Cachucho | 650 | 44.1057 | -5.1505 |
| ***Phakellia pusilla* sp. nov.** |  |  |  |  |  |
| i223 | W Mediterranean | Balearic Islands | 240 | 38.9613 | 2.0163 |
| i394_2 | W Mediterranean | Balearic Islands | 315 | 38.7828 | 2.5183 |
| i405 | W Mediterranean | Balearic Islands | 151 | 38.7062 | 3.1505 |
| i417 | W Mediterranean | Balearic Islands: Emile Baudot Seamount | 150 | 38.7327 | 2.4817 |
| i427 | W Mediterranean | Balearic Islands | 293 | 38.9668 | 2.1963 |
| i629_1_1 | W Mediterranean | Balearic Islands | 214–265 | 38.7875 | 1.7838 |
| i712 | W Mediterranean | Balearic Islands: Emile Baudot Seamount | 150 | 38.7050 | 2.4908 |
| i713_1 | W Mediterranean | Balearic Islands | 147 | 38.7050 | 2.4908 |
| i713_3 | W Mediterranean | Balearic Islands | 147 | 38.7050 | 2.4908 |
| i731 | W Mediterranean | Balearic Islands | 280–306 | 38.7060 | 2.5123 |
| i822_1 | W Mediterranean | Balearic Islands: Emile Baudot Seamount | 132 | 38.7408 | 2.4873 |
| 1170 | NE Atlantic | Canary Islands | 284 | 28.0602 | -14.2843 |
| DR7_244 | NE Atlantic | Cantabrian Sea: El Cachucho | 660 | 44.1034 | -5.1510 |
| DR14_651 | NE Atlantic | Cantabrian Sea: Avilés Canyon System | 340 | 43.8680 | -6.1425 |
| DR14_653 | NE Atlantic | Cantabrian Sea: Avilés Canyon System | 340 | 43.8680 | -6.1425 |
| ***Phakellia ventilabrum*** |  |  |  |  |  |
| M295 | NE Atlantic | Cantabrian Sea | 75 | 43.4515 | -4.2411 |
| i661 | W Mediterranean | Balearic Islands | 511 | 38.7125 | 2.4610 |
| i217 | W Mediterranean | Balearic Islands | 151–230 | 38.9598 | 2.0132 |
| i801 | W Mediterranean | Balearic Islands: Ses Olives Seamount | 320 | 38.9494 | 2.0136 |

### DNA extraction and sequencing

DNA was extracted from the subsampled tissue using the DNeasy Blood and Tissue Extraction kit (Qiagen, Hilden, Germany) and SpeedTools DNA extraction kit (Biotools, Madrid, Spain) following the manufacturer’s protocol, except for the cell lysis time, that was conducted overnight, and the final DNA elution step, that was done in 75 μL. The Folmer standard DNA barcode region of the mitochondrial cytochrome c oxidase subunit I (*cox1*), the C1–D2 and D3–D5 regions of the nuclear *28S* rDNA (*28S*), and the first fragment of the *18S* rRNA (*18S*) were amplified through polymerase chain reaction (PCR) for a representative set of individuals from each putative species (see Supplementary Table 1). Primer pairs used for the amplification of the *cox1* were the ones described in (Taboada *et al*., 2022): Pcant-COIF (5′-TTTGCAGGGATGATCGGAAC-3′) and Pcant-COIR (5′-CCCGGGGCCCTCATATTTAA-3′) for *Phakellia* aff. *robusta* sensu Díaz et al., 2025 and Prob-COIF (5’-GCGGGTATGATAGGAACAGC-3’) and Prob-COIR (5’-ACCCGGCGCTCTCATATTTA-3’) for *Phakellia robusta*. The PCR program for all *cox1* pairs of primers was 94 ◦C/5 min –(94 ◦ C/30 s–58 ◦C/30 s–72 ◦C/30 s) x 35 cycles – 72 ◦C/10 min. Primer pairs used for the amplification of the *28S* C1-D2 were C1′ASTR (5′–ACC CGC TGA ACT TAA GCA T–3′) and D2 (5′–TCC GTG TTT CAA GAC GGG–3′) (Vân Le *et al*., 1993; Cárdenas *et al*., 2011) under the PCR program 94 ◦C/5 min –(94 ◦C/15 s–56 ◦C/15 s–72 ◦C/15 s) x 30 cycles – 72 ◦ C/7 min. Primer pairs used for the amplification of the *28S* D3-D5 were 830F (5’ - CATCCGACCCGTCTTGAA - 3’) and 1520R (5’ -GCTAGTTGATTCGGCAGGTG - 3’) (Morrow *et al*., 2012) under the PCR program 94 ◦C/5 min –(94 ◦C/15 s–56 ◦C/15 s–72 ◦C/15 s) x 30 cycles – 72 ◦C/7 min. Primer pairs used for the amplification of *18S* first fragment were SP18aF (5’ - CCTGCCAGTAGTCATATGCTT - 3’) and 600R (5’ -CGAGCTTTTTAACTGCAA - 3’ (Redmond *et al*., 2007, 2013) under de PCR program 94 ◦ C/5 min –(94 ◦C/15 s–46 ◦C/15 s–72 ◦C/15 s) x 37 cycles – 72 ◦C/7 min. Amplifications were conducted in 17 μL reaction using 8 μL of VWR REDTaq DNA Polymerase 2X Master Mix (VWR, USA), 6 μL of Ultra-Pure Water, 1 μL of the forward and reverse primers and 1 μL of the DNA template. PCR products, stained with GelRed® (Biotium, USA), were visualized in a 1.5% agarose gel electrophoresis, run at 90 V for 40 min. Amplification products were purified with ExoSAP-IT (Thermo Fisher Scientific) and sequenced using the forward and reverse primers described above by MacroGen Ltd. (South Korea).

### Sequence curation and detection of introns

Consensus sequences for *cox1*, *28S* and *18S* were obtained by assembling forward and reverse DNA strands using the software *Geneious Prime* v.11.0.24+8 (Kearse *et al*., 2012). The new sequences generated were deposited in GenBank (http://www.ncbi.nlm.nih.gov/genbank/) (See Supplementary Table 1). *Phakellia robusta* is known to host a group I intron in the *cox1* (Cranston *et al*., 2021). This intron, inserted at position 387 of the *cox1* gene (based on the *Amphimedon queenslandica* complete *cox1*), can be recovered using PCR-based DNA barcoding with Prob-COIF/Prob-COIR primers. To detect whether our sequences had this intron, we aligned the newly generated *cox1* sequences against those of the intron-carrying species including *P. robusta* (GenBank = OZ183513) and *Stupenda singularis* (KP877498), as well as the intron-less reference *Amphimedon queenslandica* (ABI48988) using *MAFFT* v.7 (Katoh *et al*., 2018) with the LINS-I algorithm. After manual curation of the alignment, we selected regions that loosely aligned with intronic regions and evaluated whether they could be attributed to any extant intron group with *RNAweasel* v.1.10 (https://megasun.bch.umontreal.ca/apps/rnaweasel/). We tested for both group I and group II introns using the mold mitochondrion genetic code (Translation Table = 4). A clean intron sequence was obtained only from individual i712 (Table 1); Once intronic sequences were validated, we trimmed them and merged the exonic regions, capturing the complete orthologous *cox1* sequence.

### Phylogenetic analyses

The newly generated intron-less *cox1* (41 sequences), *28S* (C1-D2) (35 sequences), *28S* (D3-D5) (8 sequences) and *18S* (first fragment) (12 sequences) datasets were combined with published sequences retrieved from Genbank from closely related species from the order Bubarida, together with representatives of the more distantly related orders Axinellida Lévi, 1953 and Vilesida Díaz & Cárdenas, 2025. *Scopalina ruetzleri* (Wiedenmayer, 1977) was used as the outgroup in all datasets. The final datasets comprised, respectively, 85 *cox1*, 57 *28S* (C1-D2) sequences, 14 *28S* (D3-D5) and 23 *18S* (accession numbers on Figures 8–9; Supplementary Table 1). All datasets were aligned separately using *MAFFT*, and the distal parts of the sequences corresponding to primers were removed. *Gblocks* v.0.91b (Castresana, 2000) was run using default parameters for the non-coding gene *28S* to remove misaligned regions. Finally, the gene alignments lengths were: 440 bp for *cox1* (Supplementary File 1), 828 bp for *28S* (C1-D2) (Supplementary File 2), 569 bp for *28S* (D3-D5) (Supplementary File 3) and 537 bp for *18S* (Supplementary File 4). The evolutionary models to build the phylogeny were selected for each marker using the Akaike Information Criterion (AIC) in *JModeltest* v.2.1.7 (Darriba *et al*., 2012). The best-fit models were HKY + I + G for *cox1*, GTR + I + G for *28S* (C1–D2), GTR + I for *28S* (D3–D5), and K80 + I for *18S*, based on 85, 57, 14, and 23 sequences, respectively.

The alignment of each marker was run separately in *raxmlGUI* v.2.0.14 (Edler *et al*., 2021) to allow the nuclear and mitochondrial datasets to be analysed independently, using their respective evolutionary models and 1,000 bootstrap iterations. We established a bootstrap (BS) threshold value of 70 to consider a node as mildly supported, and a BS threshold value of 90 to consider it strongly supported. In parallel, we ran the same alignments in *MrBayes* v3.2.7 (Ronquist *et al.,* 2012) to infer their phylogeny using a Markov chain Monte Carlo (mcmc) approach. For each gene, we used their previously found evolutionary model with a total of 10,000,000 iterations and subsampling every 1,000 iterations, considering an initial 25% burning. We established a posterior probability (PP) threshold value of 0.9 to consider a node as mildly supported, and a PP threshold value of 0.95 to consider it strongly supported. Both the raxmlGUI and the MrBayes resulting trees were visualized and edited in *iTOL* v.7.5.1 (Letunic & Bork, 2024). Although phylogenetic trees of the first fragment of *18S* (12 sequences) and the *28S* (D3-D5) were also generated, they were excluded from the main results as these markers lacked enough resolution to resolve relationships within the group (Supplementary Fig. 3). Interspecific and intraspecific mean pairwise genetic distances using the Kimura 2-parameter model (K80; Kimura, 1980) were calculated for the *cox1* and *28S* (C1-D2) alignments in *RStudio* v.4.4.1 (R Core Team (2024)) using the packages *ape* (Paradis & Schliep, 2019) and *spider* (Brown *et al*., 2012).

For the haplotype network analyses, the *cox1* and *28S* (C1-D2) sequence alignments were manually trimmed in *Geneious Prime* for 55 specimens (438 bp) for the *cox1* and 32 for the *28S* (793 bp) including newly sequenced and NCBI samples. The software *PopART* v.1.7.2 (Leigh & Bryant, 2015) was used for building haplotype networks for each gene separately, using the TCS algorithm (Clement *et al*., 2000). *Phakellia* specimens were grouped by species in both haplotype networks.

### Morphological analysis

Spicules were obtained for 28 representative individuals of *P. robusta* (7 individuals) and *Phakellia* aff. *robusta* (21 individuals) after digestion of a 5 mm sponge fragment in bleach for 24 hours. The remaining solution was washed twice with distilled water and twice with 96% EtOH. The spicules were mounted in Eukitt^TM^ (Sigma-Aldrich, St Louis, MO, USA) mounting medium, examined under an Olympus BX43 light microscope and measured using ImageJ version 1.54n (Schneider *et al*., 2012). Spicule measurements were done for each species measuring ≥ 30 spicules per spicule category per each individual, unless otherwise indicated, comprising minimum, average and maximum lengths in micrometers (μm). The skeletal organization was studied for each species preparing thick sections of the sponge tissue for a selection of two individuals using a scalpel and cleared with clove oil for 30-60 minutes (Supplementary Table 1). For SEM images, aliquots of suspended spicules of *Phakellia* aff. *robusta* were later transferred onto aluminium stubs, air dried and sputter-coated with gold to be examined in a Scanning Electron Microscope (FEI INSPECT^TM^) at the Museo Nacional de Ciencias Naturales in Madrid, Spain and at the *Serveis Científico-tècnics* at the Universitat de les Illes Balears (UIB). The terminology applied for the morphological description of the spicules follows Boury-Esnault & Rützler (1997) and Hooper & Van Soest, (2002).

Morphometric variation of spicule width was analyzed related to the collection depth of 24 representatives of *P. robusta* and *Phakellia* aff. *robusta*. from whom exact depth was available (see Supplementary Table 1). Data visualization was performed in RStudio v.4.4.1 (R Core Team (2024)) using the *ggplot2* (Wickham, 2016) and *patchwork* (Pedersen 2025) packages. To evaluate bathymetric trends, linear regression models were fitted independently for each species and geographic basin (Atlantic vs. Mediterranean) using continuous individual specimen measurements (mean width per individual against exact depth). For each model, the slope (*β*), standard error (*SE*), coefficient of determination (*R²*), and p-value (*p*) were determined to evaluate statistical significance, along with corresponding 95% confidence intervals. For graphical visualization, specimens were grouped into 50 m depth intervals to display regional specimen ranges (absolute minimum, maximum, and mean spicule widths), while regression lines and confidence bands represent the continuous models.

### Comparative material and abbreviations

Comparative material was examined from the following institutions, for which we provide their corresponding abbreviations: CPORCANT, Colección PORíferos del CANTábrico, IEO-CSIC (Gijón, Spain); MOM, Museum of Oceanography (Monaco); PC, personal collection of P. Cárdenas, Uppsala University (Sweden); UPSZMC/UPSZTY, Zoological Collection at the Museum of Evolution (Uppsala, Sweden).

### ZooBank registration

The present article and the nomenclatural acts it contains have been registered in ZooBank, the official online registry for the International Code of Zoological Nomenclature (ICZN). The Life Science Identifier (LSID) for this publication is: urn:lsid:zoobank.org:pub:8A8710D7-FC20-45A4-AC91-E64EFA4FAC22

## Results

### Systematics

Class Demospongiae Sollas, 1885

Subclass Heteroscleromorpha Cárdenas, Pérez & Boury-Esnault, 2012)

Order Bubarida Morrow and Cárdenas, 2015

Family Bubaridae Topsent, 1894

Genus *Phakellia* Bowerbank, 1862

#### *Phakellia robusta* Bowerbank, 1866 (Fig. 2; Table 2; Supplementary Fig. 1)

##### Synonymised names

*Phacellia robusta* Bowerbank,1866 (misspelling of the genus name)

**Figure 2.**
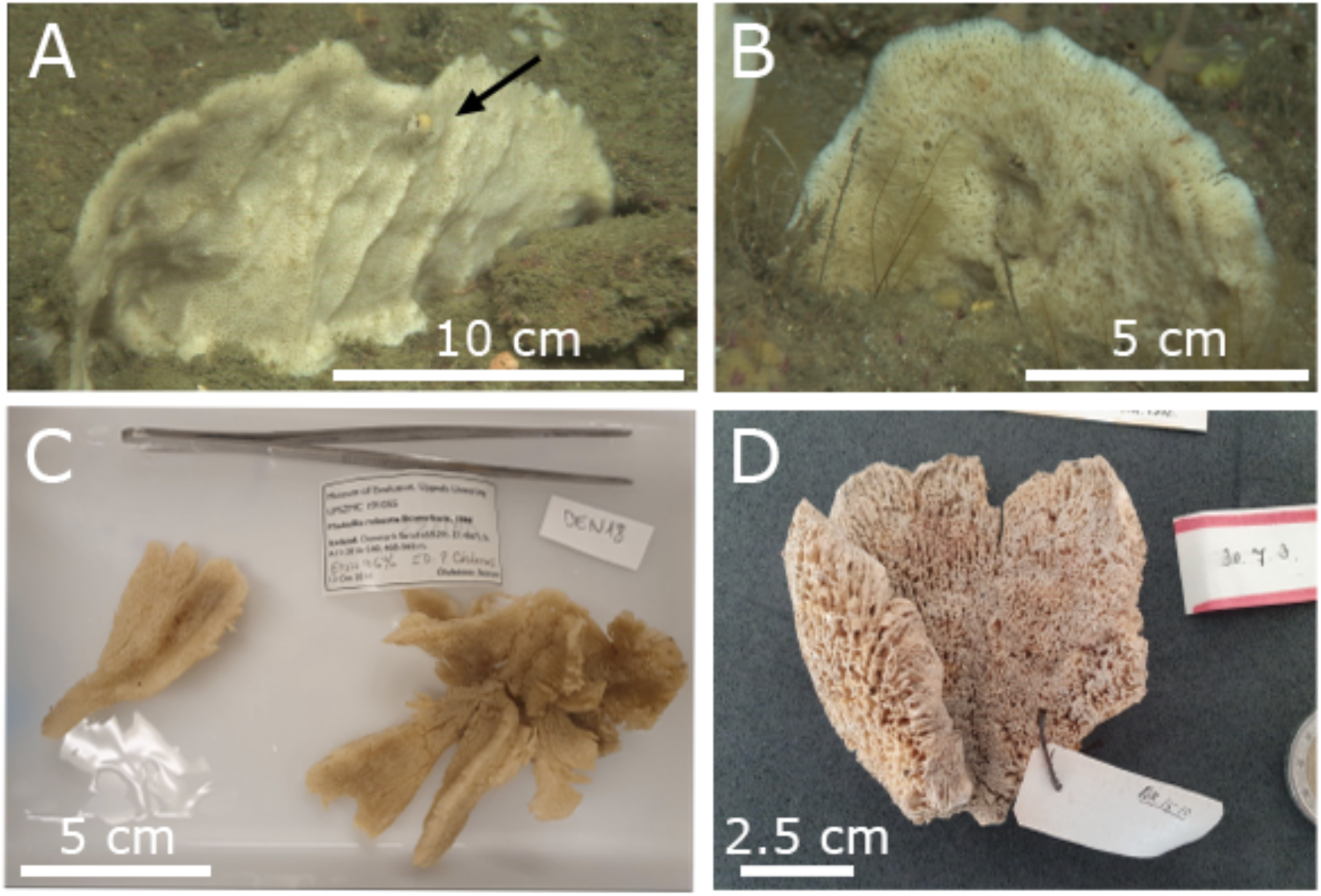
Macroscopic photographs of *Phakellia robusta* from *in situ* ROV sampling (A-B), onboard sampling (C), and museum collections (D). A: M70 from the Cantabrian Sea (NE Atlantic), 50 m. Slight striations like buttress piers are visible, and a cnidarian epibiont belonging the order Zoantharia is attached to the upper surface (marked with an arrow). B: M296 from the Cantabrian Sea (NE Atlantic), 75 m. C: UPSZMC 191055, from Denmark Strait NE Atlantic), 468-565 m. D: Type material from the Natural History Museum Zoological Collection (420).

**Table 2.** Microscopic spicule measurements of 28 individuals of *Phakellia*, including *P. robusta*, *Phakellia pallida* sp. nov. and *Phakellia pusilla* sp. nov., given as minimum-average-maximum for total length/minimum-mean-maximum for total width. Measurements for which less than 30 spicules were used are marked with *N*. Details of the measurements are given in Supplementary Table 3.

| Species/specimen | Depth (m) | Diactinal spicules | Monoactinal spicules |
| --- | --- | --- | --- |
| *Phakellia robusta* |  |  |  |
| M70 | 50 | 432 - 811 - 1,180/ 6 - 12 - 17 | 253 - 460 - 764/ 5 - 12 - 16 |
| M296 | 75 | 328 - 559 - 789/ 4 - 7 - 11 | 218 - 446 - 914/ 4 - 7 - 12 |
| M232 | 96 | 700 - 859 - 1,060/ 9 - 11 - 13 (N = 15) | 179 - 420 - 714/ 7 - 11 - 13 (N = 24) |
| 9-4_3 | 300 | 708 - 838 - 972/ 12 - 20 - 25 (N = 19) | 418 - 779 - 1,154/ 9 - 18 - 26 (N = 21) |
| 14-4_17 | 300 | 576 - 941 - 1,240/ 13 - 19 - 27 (N = 19) | 493 - 826 - 1,260/ 12 - 19 - 31 (N = 20) |
| St2_30a | 29-213 | 813 - 911 - 1022/ 11 - 18 - 30 (N = 7) | 307 - 554 - 1,126/ 13 - 18 - 24 (N = 25) |
| 420 | - | 416 - 697 - 939/ 9 - 18 - 23 (N = 21) | 249 - 563 - 1,256/ 10 - 16 - 25 (N = 29) |
| *Phakellia pallida* sp. nov. |  |  |  |
| DR10_457 | 541 | 745 - 1,177 - 1,595/ 17 - 24 - 33 (N = 20) | 229 - 352 - 591/ 11 - 15 - 22 (N = 6) |
| i347_0 | 170 | 504 - 799 - 1,169/ 7 - 11 - 16 (N = 26) | 187 - 548 - 1,160/ 4 - 9 - 13 |
| i347_2 | 170 | 551 - 688 - 902/ 5 - 8 - 10 (N = 25) | 207 - 591 - 1,299/ 5 - 8 - 15 |
| POR760 | 252 | 515 - 779 - 1,470/ 5 - 15 - 22 | 230 - 497 - 1,476/ 6 - 12 - 17 |
| POR1231 | 355 | 467 - 714 - 1,197/ 12 - 17 - 23 | 190 - 438 - 822/ 5 - 12 - 23 |
| i288 | 336-458 | 550 - 788 - 1,263/ 9 - 17 - 24 | 169 - 440 - 984/ 8 - 14 - 22 |
| i414_3 | 412 | 537 - 895 - 1,389/ 12 - 20 - 26 | 182 - 460 - 1,094/ 5 - 14 - 21 |
| DR10_455 | 541 | 518 - 724 - 1,078/ 9 - 16 - 20 (N = 27) | 196 - 400 - 633/ 9 - 13 - 17 |
| DR10_456 | 541 | 490 - 681 - 860/ 11 - 14 - 17 (N = 19) | 410 - 506 - 629/ 8 - 13 - 18 |
| DR7_253 | 660 | 630 - 796 - 1,018/ 9 - 16 - 25 (N = 27) | 358 - 709 - 1,157/ 9 - 14 - 22 (N = 26) |
| *Phakellia pusilla* sp. nov. |  |  |  |
| DR7_244 | 660 | 613 - 1,039 - 1,575/ 13 - 18 - 26 (N = 21) | 311 - 663 - 1,322/ 11 - 15 - 24 (N = 19) |
| i712 | 150 | 491 - 780 - 1,055/ 6 - 11 - 15 (N = 18) | 202 - 508 - 999/ 4 - 9 - 17 |
| i822_1 | 132 | 447 - 624 - 858/ 4 - 8 - 12 | 183 - 426 - 680/ 4 - 7 - 10 |
| i405 | 151 | 467 - 658 - 828/ 4 - 10 - 16 (N = 18) | 132 - 543 - 975/ 3 - 9 - 14 |
| i629_1_1 | 265-214 | 548 - 702 - 1,054/ 7 - 11 - 15 (N = 21) | 173 - 475 - 938/ 6 - 10 - 16 |
| i223 | 240 | 444 - 692 - 934/ 8 - 11 - 17 (N = 19) | 176 - 459 - 1,034/ 6 - 9 - 15 |
| 1170 | 284 | 571 - 695 - 844/ 9 - 13 - 19 (N = 29) | 188 - 432 - 990/ 6 - 11 - 18 |
| i427 | 293 | 627 - 900 - 1,273/ 9 - 14 - 20 (N = 25) | 214 - 642 - 1,137/ 7 - 10 - 15 |
| i394_2 | 315 | 522 - 803 - 1,273/ 12 - 17 - 21 (N = 20) | 182 - 475 - 1,056/ 7 - 12 - 19 |
| DR14_651 | 340 | 386 - 621 - 976/ 8 - 12 - 17 (N = 23) | 309 - 517 - 941/ 8 - 11 - 16 (N = 28) |
| DR14_653 | 340 | 354 - 595 - 755/ 8 - 12 - 15 (N = 26) | 163 - 405 - 605/ 5 - 9 - 14 (N = 29) |

##### Material examined

###### Holotype

**420** field no. 1877.5.21.420 NHMUK TYPE BK. 420 240 a SECTION; NHMUK 1930 7_3_187 ZOO 2025 234 ZIZ t; NHMUK 1930 7_3_188_ZOO_2025_234 ZIZ t. Shetland, United Kingdom, collected in 1864 by Gwyn Jeffreys and determined by Bowerbank, collection NHM 1877.5.21.420 (Holotype of the species). Named in the manuscript as 420.

###### North-East Atlantic

UPSZMC 193017, field no. **27-1_1**, St. 27, 14 June 2007, 292 m, Røst reef (Nordland, Norway), BT, Polarstern ARK XII/1a 2007, coll. Paco Cárdenas, det. Paco Cárdenas; UPSZMC 193014, field no. **14-4_14**, St. 14, 14 June 2007, 300 m, Røst reef (Nordland, Norway), BT, Polarstern ARK XII/1a 2007, coll. Paco Cárdenas, det. Paco Cárdenas; UPSZMC 193013, field no. **14-4_13**, St. 14, 14 June 2007, 300 m, Røst reef (Nordland, Norway), BT, Polarstern ARK XII/1a 2007, coll. Paco Cárdenas, det. Paco Cárdenas; UPSZMC 193015, field no. **14-4_17**, St. 14, 14 June 2007, 300 m, Røst reef (Nordland, Norway), BT, Polarstern ARK XII/1a 2007, coll. Paco Cárdenas, det. Paco Cárdenas; UPSZMC 193016, field no. **14-4_19**, St. 14, 14 June 2007, 300 m, Røst reef (Nordland, Norway), BT, Polarstern ARK XII/1a 2007, coll. Paco Cárdenas, det. Paco Cárdenas; UPSZMC 190312, field no. **9-4_3**, St. 14, 14 June 2007, 300 m, Røst reef (Nordland, Norway), BT, Polarstern ARK XII/1a 2007, coll. Paco Cárdenas, det. Paco Cárdenas; UPSZMC 191055, field no. **191055**, 13 October 2016, 468-565 m, Denmark strait (Iceland), BT,, Iceland A11-2016, coll. Steinunn Ólafsdóttir, det. Paco Cárdenas; field no., ST2_30a, St. 2, 12 September 2016, 29-213 m, Korsfjorden (Vestland, Norway), BT, R/V *Hans Brattstrøm*, coll. MNCN, det. Javier Cristobo; MNCN XXX, field no. **M323**, St. 13, 22 of May 2022, 96 m, Cantabrian Sea (Cantabria, Spain), ROV, CIRCAROCK-0522, coll. Ana de la Torriente, det. Carlota Gracia and Julio Diaz; MNCN XXX, field no. **M296**, St. 19, 22 of May 2022, 75 m, Cantabrian Sea (Cantabria, Spain), ROV, CIRCAROCK-0522, coll. Ana de la Torriente, det. Carlota Gracia and Julio Diaz; MNCN XXX, field no. **M70**, St. 11, 22 of May 2022, 50 m, Cantabrian Sea (Cantabria, Spain), ROV, CIRCAROCK-0522, coll. Ana de la Torriente, det. Carlota Gracia and Julio Diaz; MNCN XXX, field no. **ST6_A**, St. 6, 12 September 2016, 29-213 m, Korsfjorden (Vestland, Norway), BT, R/V *Hans Brattstrøm*, coll. MNCN, det. Javier Cristobo; MNCN XXX, field no. **KHR13**, St. 2, August 2013, 103 m, Kerry Head Reefs (County Kerry, Ireland), AD, R/V *Celtic Voyager*, det. Christine Morrow.

##### Macroscopical description (Figs. 2, 3A-C)

Lamellar to fan-shaped sponge, arranged in a semicircle, < 20 cm in height and < 30 cm in horizontal expansion. Lamellae appear as slightly curved walls, usually with some ascending striations running parallel to the shape of the wall, sometimes not reaching the very edge of the sponge (Figs. 2A-D). In some cases, lamellae can appear folded, but never fusing the ends in a cup-shaped form (Fig. 2C). In most cases, lamellae ending in a discrete semicircular attachment basis (Fig. 3A). Thickness of the sponge varies, being thicker at the base (4 - 6 mm) and thinner toward the edge (1-5 mm), where the wall becomes gradually thinner along the entire semicircular edge. Surface of lamellae smooth and flexible. Skin appears slightly hispid, with projections of the spicules at edges of the sponge. Rounded uniporal pores present in both sides of the lamella, organized following longitudinal lines across the sponge (Figs. 2B, 3B-C). Cream-colored in alive specimens; cream to beige after ethanol fixation. Mesophotic specimens appear to have a slightly darker coloration.

**Figure 3.**
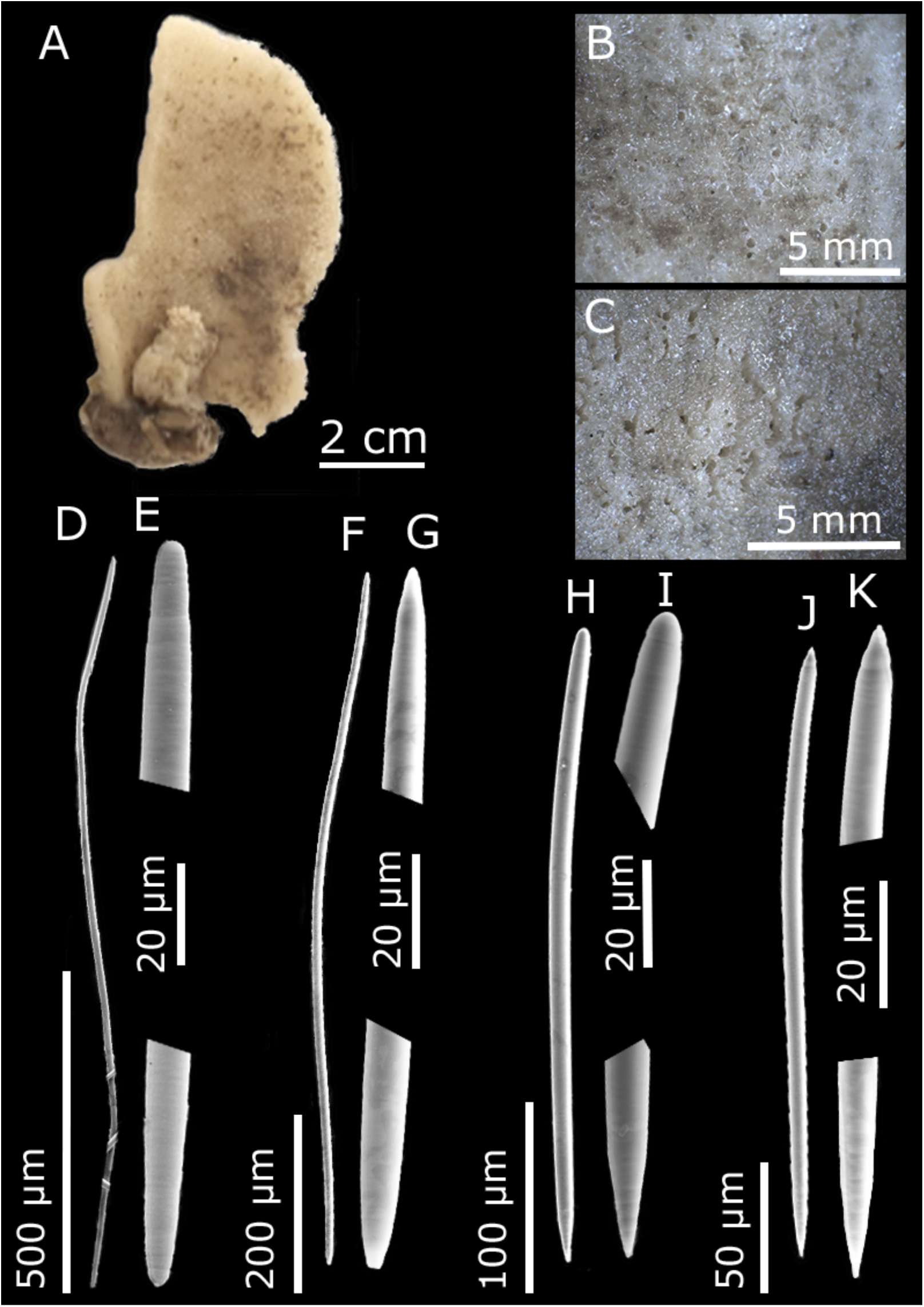
*Phakellia robusta* A: Habitus of the sample 14-4_17 from Røst Reef (NE Atlantic), 300 m showing the concave side of the lamellae. B-C: Concave and convex side (respectively) detail of the lamellae showing the uniporal pores of 14-4_17. D-K: TEM spicule images of individual M232. D-G: Diactinal sinuous spicules (D) with round ends (E) and oxeas (F) with sharp ends (G). Monactinal spicules including stylotas (H) with one round end and one sharp (I) and oxeotas (J), with one slightly pointed end and the other acutely pointed (M).

##### *Skeletal arrangement* (Supplementary Fig. 1)

The skeleton is organized in a plumoreticular arrangement, with a slender axial column giving rise to radial spicule tracts that are interconnected by large, sinuous interstitial spicules, forming a reticulated network. Towards the edges, numerous robust, non-sinuous spicules extend horizontally outwards from the radial tracts and protrude through the surface, producing a hispidated appearance.

###### Spicules

Monactinal, with unequal ends, tip always sharp, whereas head varies from rounded (stylota, Supplementary Fig. 1A) to sharp (oxeota, Supplementary Fig. 1B). Transitional morphologies present, showing telescoping of the head, which may progress to an oxeota morphology. Stylota spicules are slightly bent proximal to the head, sometimes slightly sinuous. Measuring 249 - <u>563</u> - 1,256/ 10 - <u>16</u> - 25 µm (range from the holotype).

Diactinal, with equal ends, oxeotas appearing more frequently than strongyles, with intermediate forms in between (Supplementary Fig. 1A, C, D). Irregularly curved or flexuous. Measuring 416 - <u>697</u> - 939/ 9 - <u>18</u> - 23 µm (range from the holotype).

##### *Genetics* (Figs. 9–10; Supplementary Fig. 3A)

*cox1* sequences were obtained removing the intronic region for samples 9-4_3; 14-4_13; 14-4_14; 14-4_17; 14-4_19; 27-1_1 and 191055 while no intron removal was required for the resequenced specimens (S15-13, S15-13-1, and S15-13-2) due to its absence. *28S* (C1-D2) sequences were obtained for samples ST2_30A; M232; M70 and 191055. These sequences were used to construct the phylogeny. *28S* (D3-D5) was extracted for M232 and M296, although these sequences were not included in the phylogenetic analyses because they showed little to no variation among the species studied and therefore provided no phylogenetically informative characters (Supplementary Fig. 3A).

##### Ecology and distribution

*Phakellia robusta* is present in different parts of the North-East Atlantic. The deeper samples were found in Røst reef (Nordland, Norway) ranging from 292-300 m and in the Denmark Strait (Iceland) at 468-565 m. The Cantabrian Sea samples were found shallower in the mesophotic zone ranging from 50-96 m. We cannot determine the exact depth for samples from Korsfjorden as only the start and end points of the dredging operation are available (29-213 m).

This species confers an important role in the ecosystem, forming part of the sponge grounds together with other sponges as *P. ventilabrum* or *Geodia* spp. (Sánchez *et al*., 2017). Sometimes coral epibionts of the order Zoantharia can be found on the surface of the sponge (Fig. 2A). The presence of this species in the Mediterranean should be revised, as previous records may belong to *P. pallida* sp. nov. and *P. pusilla* sp. nov. (see below).

#### *Phakellia pallida* sp. nov. Gracia-Sancha, Riesgo, Taboada & Díaz (Figures 4–5; Table 2)

*ZooBank LSID*: urn:lsid:zoobank.org:act:F2EA426B-4C18-4FAB-90E8-3A8D9DA01B9D

**Figure 4.**
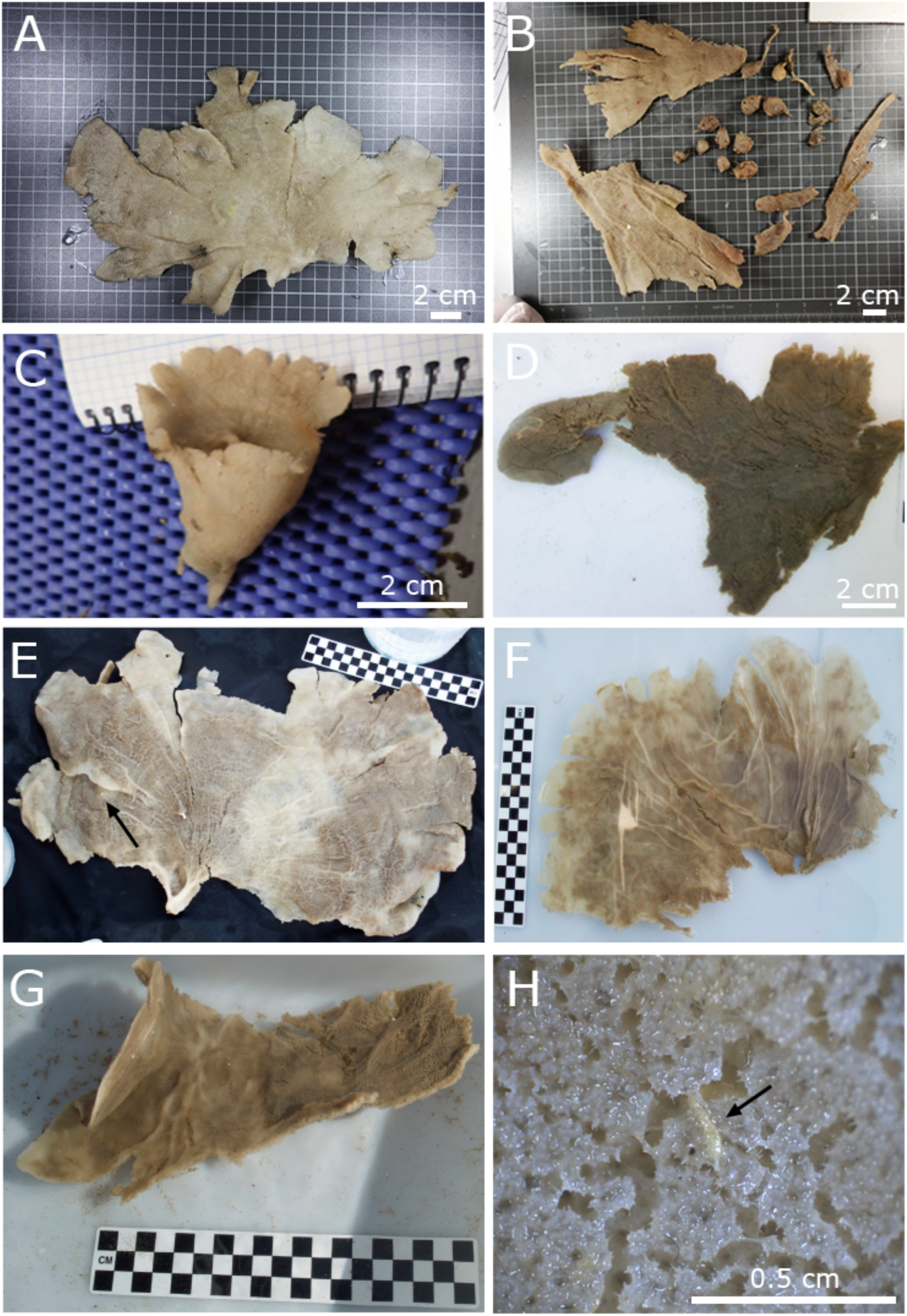
Specimens of *Phakellia pallida* sp. nov. photographed after collection. A-C. Balearic islands (W Mediterranean): A: Paratype POR462, 355 m. B: Paratype POR760, 252 m. C: POR1231, cup-shaped, 355 m. D: Avilés Canyon System (Cantabrian Sea, NE Atlantic), DR14_652,m 340 m. E-G: El Cachucho (Cantabrian Sea, NE Atlantic): E: DR10_455, 541 m. Arrow indicates an empty circle where the sponge epibiont *Craniella* sp. was living. F: DR10_456, 541 m. G: DR10_464, 541 m. H: Detail of POR760, showing a worm of the genus *Haplosyllis* (arrow) and the traces it left on the specimen.

**Figure 5.**
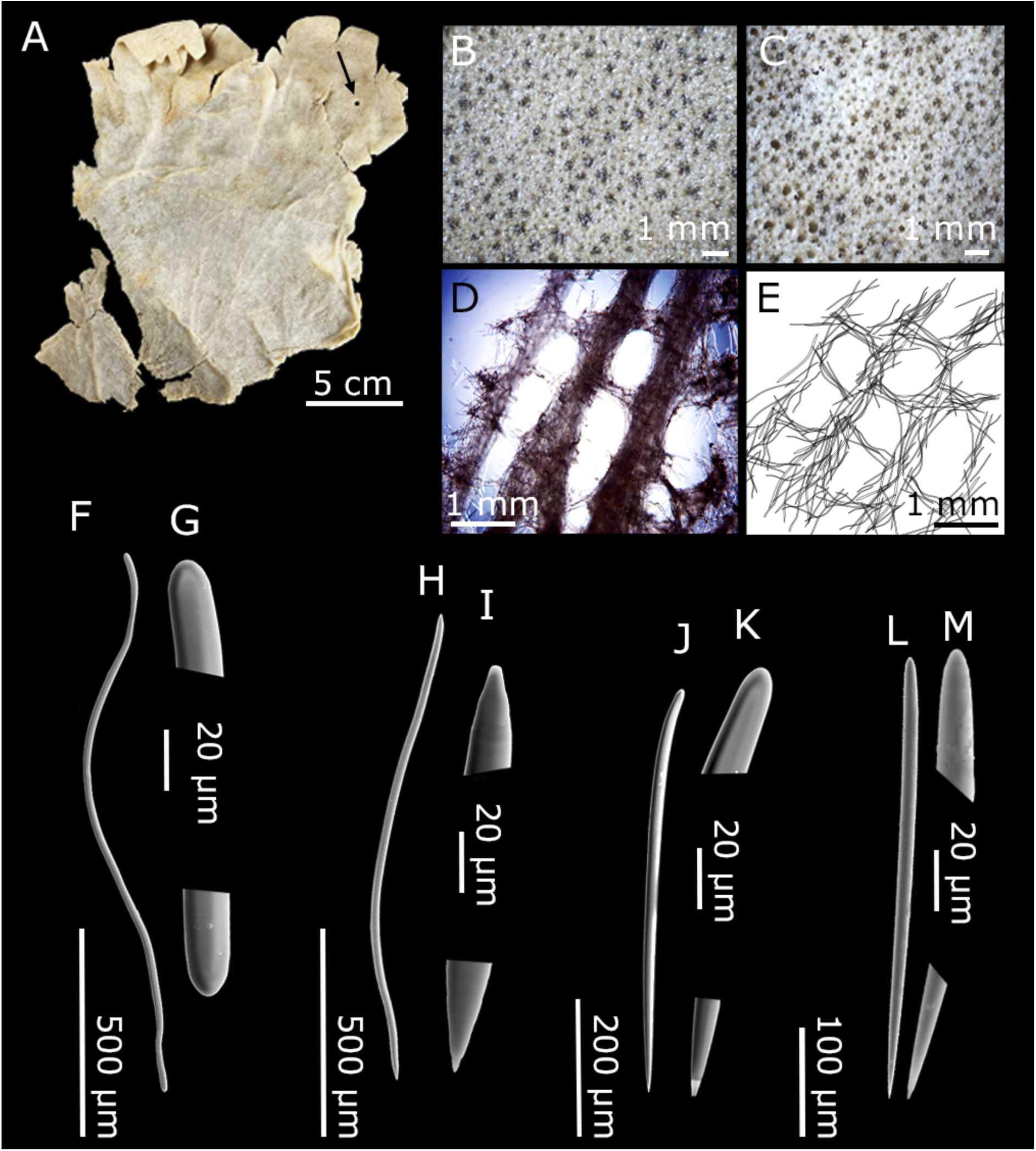
*Phakellia pallida* sp. nov. A: Habitus of the holotype DR10_457 from El Cachucho (Cantabrian Sea, NE Atlantic) showing the concave side of the lamellae. Arrow indicates an empty circle where the sponge epibiont *Craniella* sp. was living. B-C: Concave and convex side (respectively) detail of the lamellae showing the uniporal pores of DR10_457. D: Optic microscope image of the tangential section of the skeleton of POR760. E: Schematic illustration of the tangential section of the skeleton of POR760. F-M: TEM spicule images. F-I: Diactinal sinuous spicules of the holotype DR10_457 including strongyles (F) with round ends (G) and oxeas (H) with sharp ends (I). Monactinal spicules of the holotype DR10_457 including stylotas (J) with one round end and one sharp (K) and oxeotas (L), with one rounded end and with a slightly pointed one (M).

##### Synonymised names

*Phakellia robusta* Díaz *et al.,* (2021)

*Phakellia hirondellei* Ríos *et al.,* (2020), Taboada *et al.,* (2022)

*Phakellia* aff. *robusta* Díaz *et al.,* (2025)

##### Material examined

###### Holotype

MNCN XXX, field no. **DR10_457**, St. 10, El Cachucho, Cantabrian Sea, 541 m, RD, 17 June 2017, SponGES_0617, coll. P. Ríos and J. Cristobo.

###### Paratypes

MNCN XXX, field no. **POR462**, St, 212, Balearic Islands, Mediterranean Sea, 355 m, GOC-73, 13 of May 2021, Campaña MEDITS-0521, coll. J. Díaz; CFM-IEOMA-7374/POR760, field no. **POR760**, St, 224, Balearic Islands, Mediterranean Sea, 252 m, GOC-73, 24 of June 2019, Campaña MEDITS-0519, coll. J. Díaz; MNCN XXX, field no. **DR10_458**, St, 10, El Cachucho, Cantabrian Sea, 541 m, RD, 17 of June 2017, SponGES_0617, coll. P. Ríos and J. Cristobo.

###### Not part of the type series

UPSZMC 193824, field no. **ES817**, St, 20, Balearic Islands, Mediterranean Sea, 211 m, BT, 10 of June 2023, CIRCALEBA-0523, coll. J. Díaz; MNCN XXX, field no. **POR267**, Balearic Islands, Mediterranean Sea, 248 m, GOC-73, 8 of June 2017, Campaña MEDITS-0517, coll. J. Díaz; MNCN XXX, field no. **POR1050**, St, 91, Balearic Islands, Mediterranean Sea, 674 m, GOC-73, 20 of June 2020, Campaña MEDITS-0520, coll. J. Díaz; MNCN XXX, field no. **POR1231**, St, 232, Balearic Islands, Mediterranean Sea, 355 m, GOC-73, 22 of June 2021, Campaña MEDITS-0521, coll. J. Díaz; MNCN XXX, field no. **i288**, St, 97, Balearic Islands, Mediterranean Sea, 336-458 m, BT, 19 of October 2019, INTEMARES-A22B-1019, coll. J. Díaz; MNCN XXX, field no. **i347_0**, St, 135, Balearic Islands, Mediterranean Sea, 170 m, BT, 25 of October 2019, INTEMARES-A22B-1019, coll. J. Díaz; CFM-IEOMA-7367/i347_2, field no. **i347_2**, St, 135, Balearic Islands, Mediterranean Sea, 170 m, BT, 25 of October 2019, INTEMARES-A22B-1019, coll. J. Díaz; CFM-IEOMA-7369/i409, field no. **i409_1**, St, 167, Balearic Islands, Mediterranean Sea, 151 m, BT, 28 of October 2019, INTEMARES-A22B-1019, coll. J. Díaz; MNCN XXX, field no. **i414_3**, St, 175, Balearic Islands, Mediterranean Sea, 412 m, BT, 29 of October 2019, INTEMARES-A22B-1019, coll. J. Díaz; MNCN XXX, field no. **i713_4**, Balearic Islands, Mediterranean Sea, 147 m, BT, 26 of July 2020, INTEMARES-A22B-0720, coll. J. Díaz; MNCN XXX, field no. **i713_5**, Balearic Islands, Mediterranean Sea, 147 m, BT, 26 of July 2020, INTEMARES-A22B-0720, coll. J. Díaz; MNCN XXX, field no. **DR7_253**, St, 7, El Cachucho, Cantabrian Sea, 660 m, RD, 15 June 2017, SponGES_0617, coll. P. Ríos and J. Cristobo; MNCN XXX, field no. **DR9_400**, St, 9, El Cachucho, Cantabrian Sea, 660 m, RD, 16 June 2017, SponGES_0617, coll. P. Ríos and J. Cristobo; MNCN XXX, field no. **DR10_455**, St, 10, El Cachucho, Cantabrian Sea, 541 m, RD, 17 June 2017, SponGES_0617, coll. P. Ríos and J. Cristobo; MNCN XXX, field no. **DR10_456**, St, 10, El Cachucho, Cantabrian Sea, 541 m, RD, 17 June 2017, SponGES_0617, coll. P. Ríos and J. Cristobo; MNCN XXX, field no. **DR10_459**, St, 10, El Cachucho, Cantabrian Sea, 541 m, RD, 17 June 2017, SponGES_0617, coll. P. Ríos and J. Cristobo; MNCN XXX, field no. **DR10_464**, St, 10, El Cachucho, Cantabrian Sea, 541 m, RD, 17 June 2017, SponGES_0617, coll. P. Ríos and J. Cristobo; MNCN XXX, field no. **DR10_466**, St, 10, El Cachucho, Cantabrian Sea, 541 m, RD, 17 June 2017, SponGES_0617, coll. P. Ríos and J. Cristobo; MNCN XXX, field no. **DR10-498**, St, 10, El Cachucho, Cantabrian Sea, 541 m, RD, 17 June 2017, SponGES_0617, coll. P. Ríos and J. Cristobo; MNCN XXX, field no. **DR14_652,** St, 14, Avilés Canyon System, Cantabrian Sea, 340 m, RD, 21 June 2017, SponGES_0617, coll. P. Ríos and J. Cristobo; MNCN XXX, field no. **DR15_857b**, St, 15, El Cachucho, Cantabrian Sea, 650 m, RD, 23 June 2017, SponGES_0617, coll. P. Ríos and J. Cristobo; MNCN XXX, field no. **DR15_857c**, St, 15, El Cachucho, Cantabrian Sea, 650 m, RD, 23 June 2017, SponGES_0617, coll. P. Ríos and J. Cristobo; MNCN XXX, field no. **DR15_976**, St, 15, El Cachucho, Cantabrian Sea, 650 m, RD, 23 June 2017, SponGES_0617, coll. P. Ríos and J. Cristobo. All of the SponGES specimens originally identified as *Phakellia hirondellei* in Taboada et al., 2022.

MOM: **MON33** (working code assigned by the author), museum code MOM 041288, st. 1304, Josephine Bank, Azores, North-East Atlantic, 208 m, RD, 27 July 1902, dried individual first identified as *Phakellia hirondellei* by Topsent (1928) and as *Phakellia* aff. *robusta* by Díaz et al. (2025).

Note: CFM-IEOMA-7367/i347_2, CFM-IEOMA-7369/i409, CFM-IEOMA-7373/i731 and CFM-IEOMA-7374/POR760 and were reported as *P. robusta* in Díaz et al. 2021 and in Díaz et al., 2024.

##### Macroscopical description (Figs. 4, 5A)

Single lamellar to fan-shaped sponge, body usually ≤ 30 cm in height and ≤ 40 cm in horizontal expansion (Fig. 4A-B). Small individuals can be found ranging from 0.5 - 7 cm in height and 0.3 - 4 cm in horizontal expansion (Fig. 4C-D). Margins irregular and slightly undulating, extending from a hard stalk anchored to the substrate (Figs. 4E-G, 5A). Lamellae broad and curved, usually with visible branching striations radiating from a central point of the stalk, and bifurcating towards the end of the lamellae, similar to veins in a leaf or the structure of a tree, not reaching the very edge of the sponge (Fig. 4C, F-G). Unusually, lamellae can appear folded in a cup-shaped form (Fig. 4C). Thickness of the sponge consistent along the body (1.15 - 3.3 mm). Consistency smooth and flexible. A concave and a convex part can be distinguished in the lamellae, with pores on both sides (87-876 µm in diameter). Grooves observed sometimes in the skin, produced by annelid Syllidae worms that live inside the sponge (Fig. 4H). Rounded uniporal pores present in both sides of the lamella, organized following longitudinal lines across the sponge, usually more visible on one side than the other (Fig. 5B-C). From cream-colored to brown in alive specimens; cream to beige after ethanol fixation. Díaz *et al.,* (2025) provide in situ photographs of specimen ES817.

##### *Skeletal arrangement* (Fig. 5D-M; Table 2; Supplementary Fig. 2)

The skeleton is organized in a plumo-reticular net supported by tracts of sinuous spicules drawing regular meshes (Fig. 5D-E). At the edges, a greater number of non-sinuous spicules can be seen extending horizontally outwards and protruding the surface, causing hispidation (Fig. 5D).

###### Spicules

Diactinal, with equal ends, oxeas (Fig. 5H-I) appearing more frequently than strongyles (Fig. 5F-G), with intermediate forms in between. Irregularly curved or flexuous. Measuring 745 - <u>1,177</u> - 1,595/ 17 - <u>24</u> - 33 µm (range from the holotype).

Monactinal, with unequal ends, tip always sharp, with head varying from rounded (stylota, Fig. 5J-K) to sharp (oxeota, Fig. 5L-M). Transitional morphologies present, showing telescoping of the head, which may progress to an oxeota morphology. In some cases, stylota forms are slightly bent proximal to the head, and may even appear being slightly sinuous. Measuring 229 - <u>352</u> - 591/ 11 - <u>15</u> - 22 µm (range from the holotype).

##### *Genetics* (Figs. 9–10; Supplementary Fig. 3)

*cox1* sequences were for samples MON33; POR267; POR462; POR760; POR405; POR1231; i288; i347_0; i347_2; i409_1; i414_3; i713_4 and i713_5 whereas *28S* (C1-D2) sequences were obtained for samples MON33; POR462; POR760; POR1050; POR1231; i347_0; i347_2; i409_1; i414_3; i713_4; i713_5; DR9_400; DR10_455; DR10_457; DR10_498; DR14_652. These sequences were used to construct the phylogeny. *28S* (D3-D5) was extracted for MON33 and DR10_457 and *18S* (first fragment) was extracted for POR760; POR1231; i347_0; i347_2; i414_3; i713_4 and DR10_457. These sequences were not included in the phylogenetic analyses because they showed little to no variation among the species studied and therefore provided no phylogenetically informative characters (Supplementary Fig. 3).

##### Ecology and distribution

*Phakellia pallida* sp. nov. is present in the Balearic Islands (W Mediterranean), the Cantabrian Sea (NE Atlantic) and in Azores (NE Atlantic). The Bathymetric range varies from 151 to 660 m. This species could potentially form part of multispecific sponge grounds in the Cantabrian Sea. However, its actual distribution and abundance in this area may be underestimated, as some specimens previously identified as *P. robusta* may correspond to this species or other closely related taxa within the *P. robusta* species complex. It occurs alongside other sponges such as *P. ventilabrum*, *P. robusta* and *Geodia barretti* Bowerbank, 1858 (Taboada *et al*., 2022; Rodríguez-Basalo *et al*., 2026). In the Mediterranean, this species normally occurs in isolated patches in rocky outcrops and vertical walls (Díaz *et al*., 2021). In some cases, annelid syllids of the genus *Haplosyllis* Langerhans, 1879 are found living inside this species, as well as sponge epibionts of the genus *Craniella* Schmidt, 1870 living on its surface (Figs. 4H, 5A).

At the Balearic archipelago, the species is commonly captured as a by-catch by bottom trawlers, particularly at mud bottoms of the upper and middle continental slope (reported as *P. robusta* in (Díaz, 2024).

###### Etymology

From latin *pallida* (pale), referring to the color of this species.

#### *Phakellia pusilla* sp. nov. Gracia-Sancha, Riesgo, Taboada & Díaz (Figures 6–7; Table 2)

*ZooBank LSID*: urn:lsid:zoobank.org:act:8AC0240A-7D12-4BAC-AD71-8976B3986273

**Figure 6.**
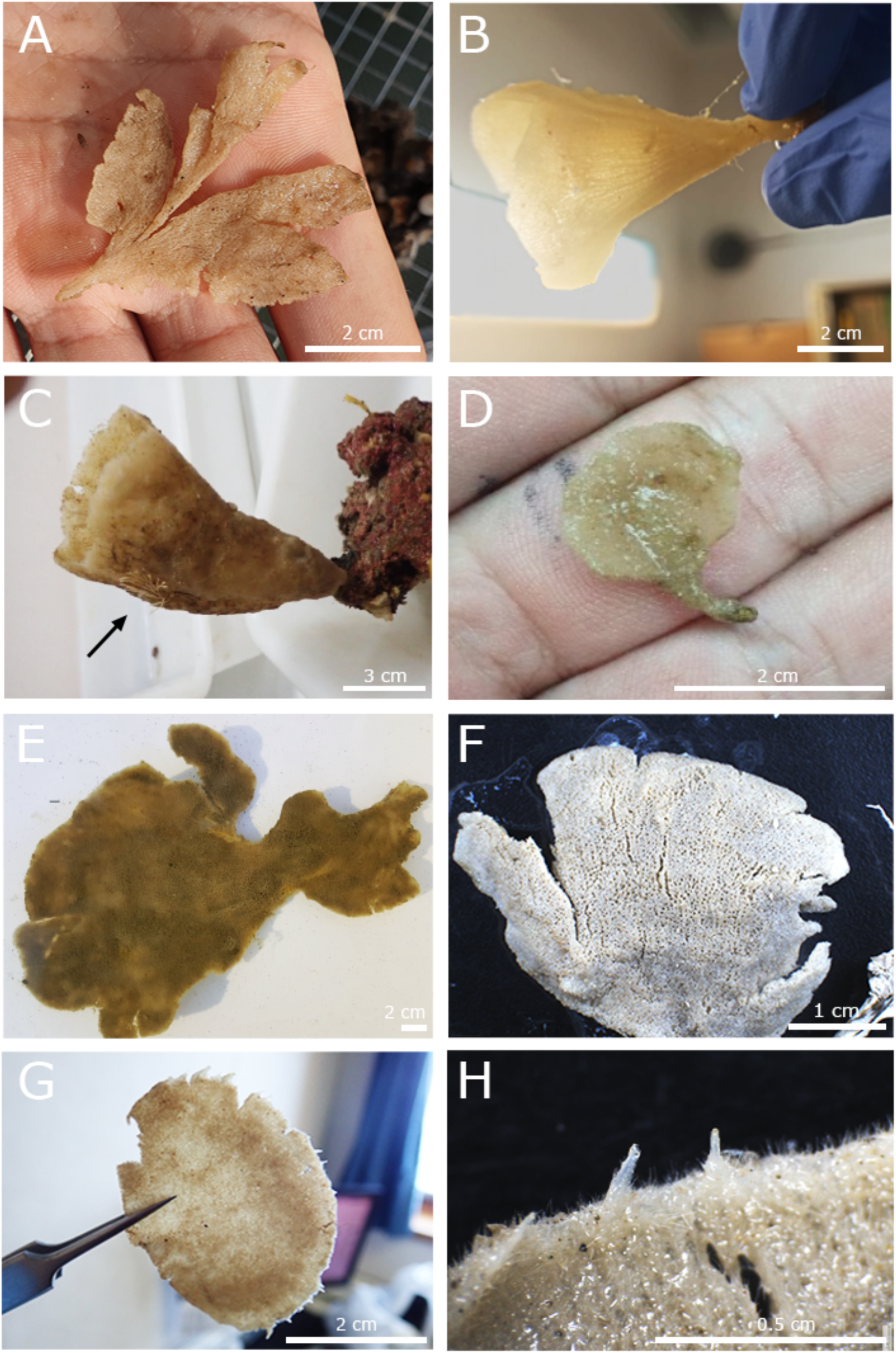
Specimens of *Phakellia pusilla* sp. nov. photographed after collection. A: Specimen i394_2 collected from the Balearic Islands (W Mediterranean) at 315 m. B: 1170, cup-shaped, from the Canary Islands (Atlantic Ocean), 284 m. C: Paratype i822_1, cup-shaped, from the Balearic Islands (W Mediterranean), 132 m. Arrow indicates the presence of a bryozoan. D: i405 collected from the Balearic Islands (W Mediterranean), 151 m. E: DR14_651, from the Avilés Canyon System (Cantabrian Sea, NE Atlantic), 340 m. F: Paratype i223, from the Balearic Islands (W Mediterranean), 240 m. G: i427 from the Balearic Islands (W Mediterranean), 293 m. H: Detail of the distal part of the body of i427 where reproductive extensions in the form of buds can be appreciated.

**Figure 7.**
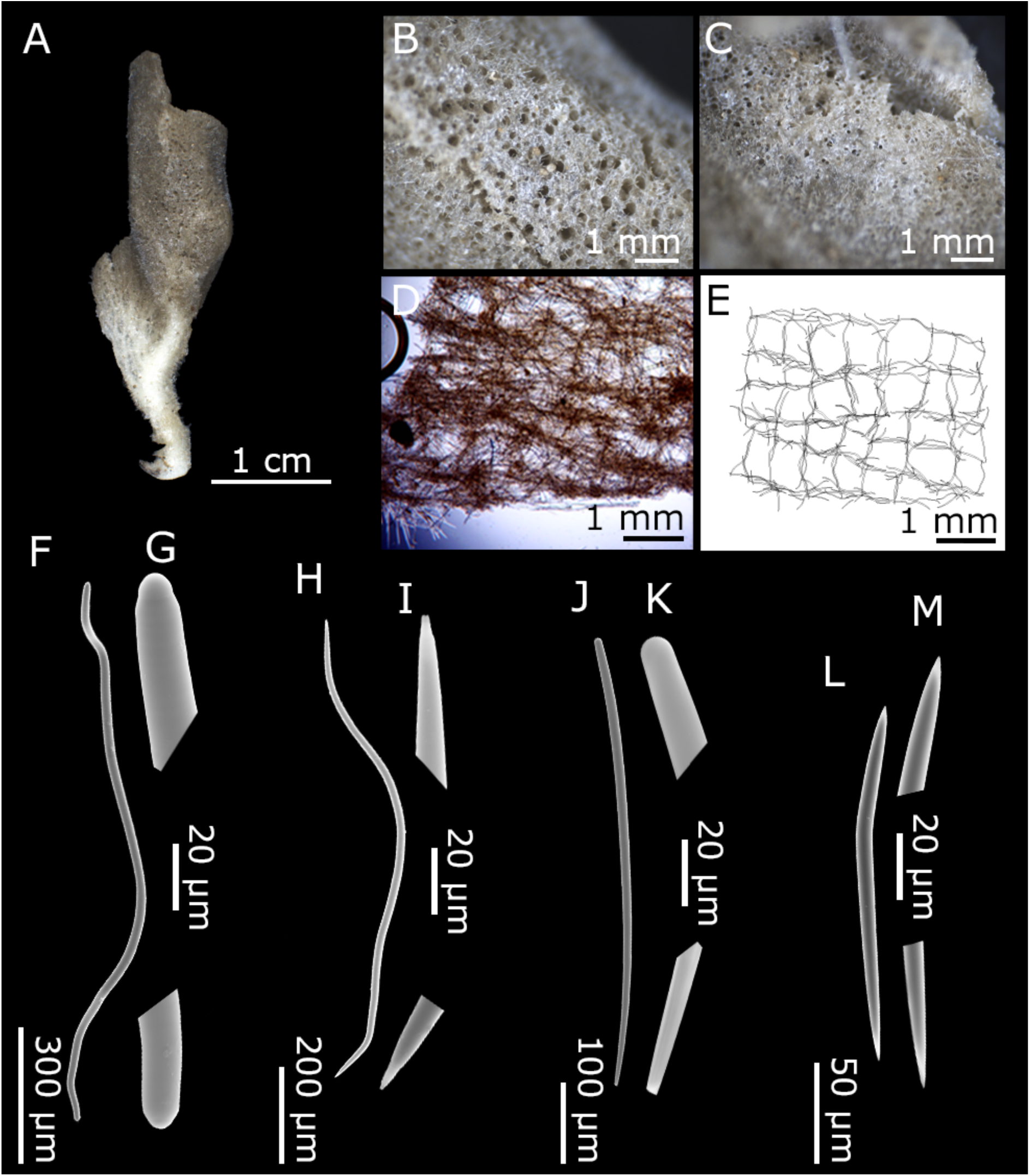
Holotype and type series of *Phakellia pusilla* sp. nov. A: Habitus of the holotype DR7_244 from El Cachucho (Cantabrian Sea, NE Atlantic) showing the convex side of the lamellae. B-C: Concave and convex side (respectively) detail of the lamellae showing the cribiporal pores of DR7_244. D: Light microscopy image of the tangential section of the skeleton of i822_1. E: Schematic illustration of the tangential section of the skeleton of i822_1. F-M: TEM spicule images. F-I: Diactinal sinuous spicules of the holotype DR7_244 including strongyles (F) with round ends (G) and oxeas (H) with sharp ends (I). Monactinal spicules of the holotype DR7_244 including stylotas (J) with one round end and one sharp (K) and oxeotas (L), with one rounded end and with a slightly pointed one (M).

##### Synonymised names

*Phakellia ventilabrum* Díaz *et al.,* (2021)

*Phakellia robusta* Díaz *et al.,* (2021)

*Phakellia robusta* Taboada *et al.,* (2022) referred to as a hybrid.

*Phakellia* aff. *robusta* Díaz *et al.,* (2025)

##### Material examined

###### Holotype

MNCN XXX, field no. **DR7_244**, St. 7, El Cachucho, Cantabrian Sea, 660 m, RD 15 June 2017, SponGES_0617, coll. P. Ríos and J. Cristobo.

###### Paratypes

MNCN XXX, field no. **i223**, Balearic Islands, Mediterranean Sea, 240 m, BT, 11 of October 2019, INTEMARES-A22B-1019, coll. J. Díaz;

###### Not part of the type series

MNCN XXX, field no. **i394_2**, St, 165, Balearic Islands, Mediterranean Sea, 315 m, BT, 28 of October 2019, INTEMARES-A22B-1019, coll. J. Díaz; CFM-IEOMA-7368/i405, field no. **i405** St, 167, Balearic Islands, Mediterranean Sea, 151 m, BT, 28 of October 2019, INTEMARES-A22B-1019, coll. J. Díaz; CFM-IEOMA-7371/i417 field no. **i417**, St, 177, Emile Baudot seamount, Balearic Islands, Mediterranean Sea, 150 m, BT, 29 of October 2019, INTEMARES-A22B-1019, coll. J. Díaz; MNCN XXX, field no. **i427,** Balearic Islands, Mediterranean Sea, 293 m, BT, 21 of July 2020, INTEMARES-A22B-0720, coll. J. Díaz; MNCN XXX, field no. **i629_1_1**, Balearic Islands, Mediterranean Sea, 214-265 m, BT, 24 of July 2020, INTEMARES-A22B-0720, coll. J. Díaz; UPSZMC 193828, field no. **i712**, St, 45, Emile Baudot seamount, Balearic Islands, Mediterranean Sea, 150 m, BT, 26 of July 2020, INTEMARES-A22B-0720, coll. J. Díaz; MNCN XXX, field no. **i713_1**, Balearic Islands, Mediterranean Sea, 147 m, BT, 26 of July 2020, INTEMARES-A22B-0720, coll. J. Díaz; MNCN XXX, field no. **i713_3**, Balearic Islands, Mediterranean Sea, 147 m, BT, 26 of July 2020, INTEMARES-A22B-0720, coll. J. Díaz; CFM-IEOMA-7373/i731, field no. **i731**, St, 46, Balearic Islands, Mediterranean Sea, 280-306 m, BT, 26 of July 2020, INTEMARES-A22B-0720, coll. J. Díaz; CFM-IEOMA-7376/i822_1, field no. **i822_1**, St, 23, Emile Baudot seamount, Balearic Islands, Mediterranean Sea, 133-169 m, BT, 29 of August 2020, INTEMARES-A22B-0820, coll. J. Díaz; MNCN XXX, field no. **1170,** St, 3, Canary Islands, Atlantic Ocean, 284 m, ROV, 26 of November 2021, CANZEC-2021, coll. M. González-Porto; MNCN XXX, field no. **DR14_651**, St, 14, Avilés Canyon System, Cantabrian Sea, 340 m, RD, 21 of June 2017, SponGES_0617, coll. P. Ríos and J. Cristobo; MNCN XXX, field no. **DR14_653**, St, 14, Avilés Canyon System, Cantabrian Sea, 340 m, RD, 21 of June 2017, SponGES_0617, coll. P. Ríos and J. Cristobo.

###### Phakellia ventilabrum

MNCN XXX, field no. **M295**, St, 19, Cobreces, Cantabrian Sea, 75 m, ROV, 26 of May 2022, CIRCAROCK-0522, coll. A. de la Torriente; MNCN XXX, field no. **i661,** St, 38, Balearic Islands, Mediterranean Sea, 511 m, BT, 26 of July 2020, INTEMARES-A22B-0720, coll. J. Díaz; MNCN XXX, field no. **i217**, St, 8, Balearic Islands, Mediterranean Sea, 151-230 m, BT, 11 of October 2019, INTEMARES-A22B-1019, coll. J. Díaz; MNCN XXX, field no. **i801**, St, 5-2, Ses Olives seamount, Balearic Islands, Mediterranean Sea, 320 m, BT, 24 of August 2020, INTEMARES-A22B-0820, coll. J. Díaz.

Note: CFM-IEOMA-7376/i822_1 was reported as *Phakellia ventilabrum* in Díaz et al., 2021 and Diaz et al., 2024, CFM-IEOMA-7368/i405 and CFM-IEOMA-7371/i417, CFM-IEOMA-7372/i712, were reported as *P. robusta* in Diaz et al., 2021 while i217 and i661 were reported as *Phakellia* sp. in Díaz *et al*. 2024.

##### Macroscopical description (Figs. 6, 7A-C)

Single lamellar to fan-shaped sponge, with body 2 - 7 cm in height and 1.5 - 10 cm in horizontal expansion (Fig. 6). Margins regular and extending from a stalk that is anchored to the substrate (Fig. 6A-D). Lamellae broad and flattened, folded and fused in some big individuals in a cup-shaped form, similar to that of *P. ventilabrum* (Fig. 6B-C). Thickness of the sponge relatively consistent across body, ranging from 0.7 to 1.3 mm depending on the size of the individuals, with larger specimens being thicker. Surface of lamellae smooth and flexible. A concave and a convex part can be distinguished in the lamellae, with cribiporal pores on both sides (61-586 µm in diameter) organized following longitudinal lines across the sponge with smaller pores consistently appearing on one side of lamellae (Fig. 7B-C). From cream-colored to brown in alive specimens (Fig. 6A, D, E); cream to beige after ethanol fixation (Fig. 6D, F-G; Fig. 7A). Díaz *et al.,* (2021) provide in situ photographs of specimen i822_1.

##### *Skeletal arrangement* (Fig. 7D-M; Table 2; Supplementary Fig. 2)

The skeleton is organized in a plumo-reticular net supported by tracts of sinuous spicules drawing regular meshes (Fig. 7D, E). At the edges, a greater number of non-sinuous spicules can be seen extending horizontally outwards, protruding the surface and causing hispidation (Fig. 7D).

###### Spicules

Diactinal, with equal ends, oxeas appearing more frequently than strongyles, with intermediate forms in between (Fig. 7F-I). Irregularly curved or flexuous. Measuring 613 - <u>1,039</u> - 1,575/ 13 - <u>18</u> - 26 µm (range from the holotype).

Monactinal, with unequal ends, tip always sharp, whereas head varies from rounded (stylota, Fig. 7J-K) to sharp (oxeota, Fig. 7L-M). Transitional morphologies present, showing telescoping of the head, which may progress to an oxeota morphology. In some cases, stylota forms are slightly bent proximal to the head, and may even appear being slightly sinuous. Measuring 311 - <u>663</u> - 1,322/ 11 - <u>15</u> - 24 µm (range from the holotype).

##### *Genetics* (Figs. 9–10; Supplementary Fig. 3)

The *cox1* fragment sequences were extracted for samples i223; i394_2; i405; i417; i427; i629_1_1; i712; i713_1; i347_2; i713_3; i731; i822_1; 1170; DR7_244; DR14_651 and DR14_653 in which individuals i417, i629_1_1, i712, i713_3, i822_1 presented an intronic region of 1356 bp. In addition, *28S* (C1-D2) sequences were obtained for samples i223; i394_2; i427; i712; i713_1; i731; i822_1; 1170; DR7_244 and DR14_653. These sequences were used to construct the phylogeny. Finally, *28S* (D3-D5) was extracted for i712; i822_1; DR7_244 and 1170 and *18S* (first fragment) was extracted for DR7-244. These sequences were not included in the phylogenetic analyses because they showed little to no variation among the species studied and therefore provided no phylogenetically informative characters (Supplementary Fig. 3).

##### Ecology and distribution

*Phakellia pusilla* sp. nov. is present in the Balearic Islands (W Mediterranean Sea), the Canary Islands (NE Atlantic Ocean) and the Cantabrian Sea (NE Atlantic). Its bathymetric range is located between 147 and 660 m. This species forms part of multispecific sponge grounds in the Cantabrian Sea, where individuals may reach larger sizes, although this remains uncertain since only partial specimens have been collected from this area. However, its actual distribution and abundance in this area may be underestimated, as some specimens previously identified as *P. robusta* may correspond to this species or other closely related taxa within the *P. robusta* species complex. It occurs alongside other sponges such as *P. ventilabrum* and *P. robusta* (Rodríguez-Basalo *et al*., 2026). In the Balearic Islands, *P. pusilla* sp. nov. does not appear to be found in large densities, although it was frequent in the Emile Baudot seamount upper slopes. Bryozoans have been observed as epibionts on this species (Fig. 6D). Asexual reproductive structures in the form of buds were observed in one specimen (i427) in the apical part of the body (Fig. 6F).

###### Etymology

From latin *pusilla* (feminine adjective meaning “small”), referring to the smaller size of this species compared to their sister species *P. robusta* and *P. pallida* sp. nov.

### Morphological analyses

Morphometric variations of spicule width of diactinal (Supplementary Fig. 1A) and monactinal (Supplementary Fig. 1B; Supplementary Table 2) spicules increased in general with depth in the species analyzed. For diactinal spicules, specimens of *P. robusta* from the Atlantic showed an increase from 10 µm at 50-100 m depth to 19 µm at 300-350 m (*β* = 0.041 ± 0.010, *R²* = 0.85, *p* = 0.027). In *P. pusilla* sp. nov., widths increased from 12 µm at 300-350 m to 17 µm at 600-700 m in Atlantic specimens (*β* = 0.019, *R²* = 1.00, *p* < 0.001, albeit constrained by the number of individuals (3)), and significantly from 9 µm at 100–150 m to 17 µm at 300-350 m in Mediterranean specimens (*β* = 0.034 ± 0.008, *R²* = 0.80, *p* = 0.007). Similarly, *P. pallida* sp. nov. from the Mediterranean exhibited a strong significant increase from 9 µm at 150-200 m to 20 µm at 400-450 m (*β* = 0.042 ± 0.007, *R²* = 0.91, *p* = 0.011). However, *Phakellia pallida* sp. nov. from the Atlantic showed a slight, non-significant variation in the width from 18 µm at 500-550 m to 17 µm at 650-700 m (*β* = -0.017 ± 0.051, *R²* = 0.05, *p* = 0.775).

For monactinal spicules, specimens of *P. robusta* from the Atlantic showed a significant increase from 10 µm at 50-100 m depth to 18 µm at 300-350 m (*β* = 0.037 ± 0.010, *R²* = 0.82, *p* = 0.034). In Mediterranean populations, spicule width was positively and significantly correlated with depth in both *P. pusilla* sp. nov., increasing from 9 µm at 100-150 m to 17 µm at 300-350 m (*β* = 0.018 ± 0.005, *R²* = 0.74, *p* = 0.013), and *P. pallida* sp. nov., shifting from 8 µm at 150-200 m to 12 µm at 400-450 m (*β* = 0.021 ± 0.005, *R²* = 0.86, *p* = 0.023). In contrast, Atlantic specimens did not yield statistically significant bathymetric correlations: *P. pusilla* sp. nov. increased from 10 µm at 300-350 m to 15 µm at 600-700 m without reaching significance (*β* = 0.016 ± 0.005, *R²* = 0.89, *p* = 0.212), while Atlantic *P. pallida* sp. nov. showed an apparent reduction from 13 µm at 500-550 m to 9 µm at 650-700 m that lacked statistical support (*β* = 0.003 ± 0.011, *R²* = 0.03, *p* = 0.826).

### Sequence assembly, intron identification, and phylogenetic analysis

Five individuals from Mallorca (i712, i417, i629_1_1, i713_3, i822_1) presented *cox1* sequences that were larger (>1,300 bp) than the expected ca. 600 bp for sponge barcodes, suggesting the presence of an intron. All these individuals belonged to the new species *Phakellia pusilla* sp. nov. The specimen i712 yielded a high-quality assembly for the entire 1,300 bp and was used as the reference sequence to determine position and type of the potential intron, although all of them had the same sequence structure. When producing an alignment composed by the sequences of the i712 specimen, *P. robusta*, *S. singularis*, and *A. queenslandica*, we detected that the non-*cox1* region in the specimen i712 was homologous to the intron of *P. robusta* and *S. singularis* (Supplementary Fig. 4). The region homologous to the intron was inserted in position 387 relative to *A. queenslandica*, and spanned a total of 1,356 bp (Fig. 8A). The insertion site was coincident with those of *P. robusta* and *S. singularis* introns, yet their lengths differed (1,752 and 1,044 bp, respectively) (Fig. 8B). The RNAweasel evaluation determined that the homologous region corresponded to a group IB intron (Hausner *et al*., 2014). For further analyses, in order to include these individuals in the phylogenies constructed, the intronic regions were removed from all the sequences containing introns following the procedure described in Taboada et al. (2022), and alignments were performed only on the exonic regions.

**Figure 8.**
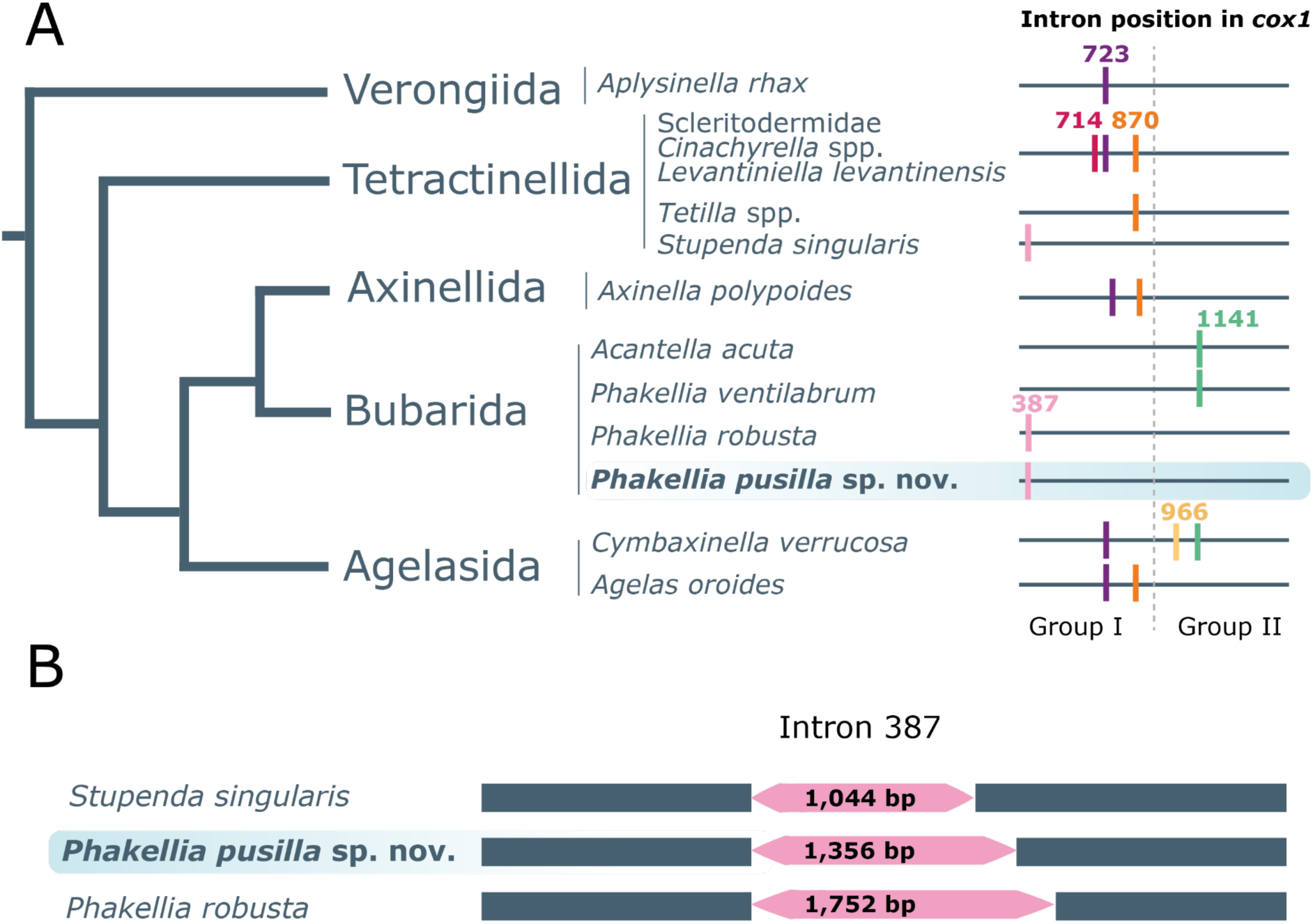
A: Reconstruction of the known mitochondrial introns in the *cox1* gene in sponges based on a simplified phylogeny. Numbers indicate the insertion position of the introns and distinguish between Group I and Group II introns. Modified from Schuster *et al.,* (2017) with findings from Galià-Camps *et al*., (2026) of the intron found in *P. ventilabrum*. the intronic position of some individuals of *Phakellia pusilla* sp. nov. (i417, i629_1_1, i712, i713_3, i822_1), belonging to group I, is highlighted in blue. B: Representation of the alignment of *cox1* fragments from sponge species containing an intronic region inserted at position 387, indicating the length variation of the intronic region.

#### Phylogenetic analyses

The Maximum-likelihood phylogenetic trees built for the *cox1* (Fig. 9) and the *28S* (Fig. 10) included representatives of three related orders, Axinellida, Bubarida and Vilesida. Axinellida and Vilesida formed a strongly supported sister clade (*cox1* and *28S* BS = 100) in both trees, and then Bubarida species clustered separately from this group in a well supported clade (*cox1* BS = 99; *28S* BS = 100). All analysed *Phakellia* specimens allocated within Bubarida, but did not form a monophyletic group, forming instead a polyphyletic group in both *cox1* and *28S* datasets (Figs. 9–10, respectively). *Phakellia ventilabrum,* the type species for *Phakellia,* produced a highly supported clade (*cox1* and *28S* BS = 99), appearing as a sister species of *Axinella cannabina* (Esper, 1794) in both trees. The remaining *Phakellia* species (*P. robusta* and *Phakellia* aff. *robusta* sensu Díaz et al., 2025) formed a clade that we called *P. robusta* species complex, that was recovered with moderate support in both the *cox1* and *28S* phylogenetic trees (Figs. 9–10). Specimens of the *P. robusta* species complex were sister to members of the genus *Cymbastela* Hooper & Bergquist, 1992 and *Dictyonella* Schmidt, 1868 for both trees (*cox1* and *28S* BS = 99). Within this clade, *P. robusta* and *Phakellia* aff. *robusta* sensu Díaz et al., 2025 formed a well-supported monophyletic group (*28S* BS = 100; *cox1* BS = 97).

**Figure 9.**
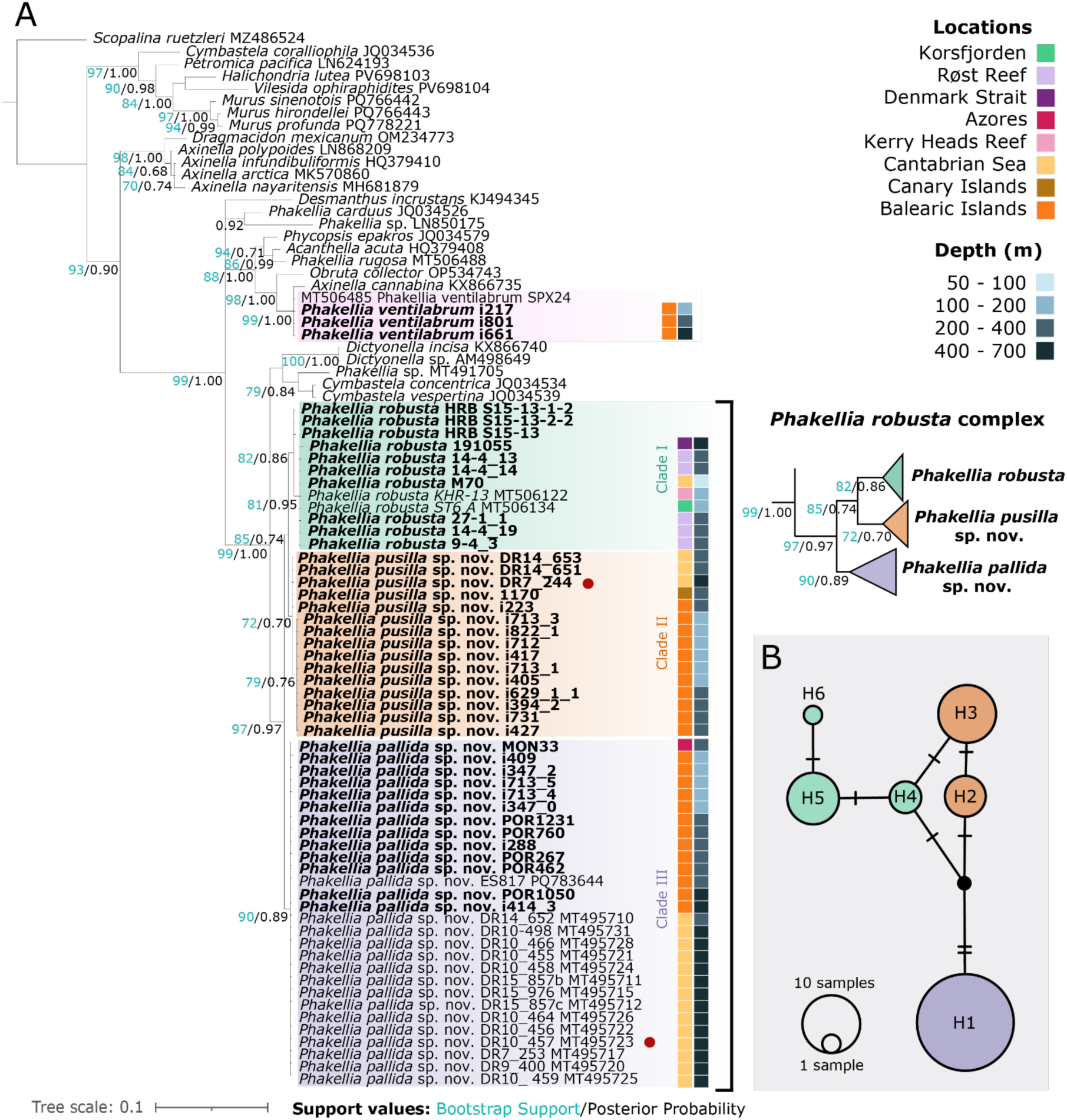
A: Bayesian 50% majority-rule consensus tree inferred with MrBayes based on the *cox1* gene. Numbers at nodes indicate Maximum Likelihood bootstrap support (1,000 replicates; green) followed by Bayesian posterior probabilities (black). Only support values with BS > 70% or PP > 0.70 are shown. Genbank accession numbers appear after each taxon name. Specimens sequenced in our study are in bold. Colored squares show both sampling locations and depth. Red dots mark the individuals designated as type specimens. A magnified view of the phylogenetic tree showing *Phakellia robusta* complex is displayed on the right. B: Haplotype network based on the *cox1* for the *Phakellia robusta* complex. Circles are proportional to the number of individuals for each haplotype. Cross lines symbolize the mutational steps and missing inferred haplotypes are in black. Each color represents a different species: *Phakellia robusta* green, *Phakellia pusilla* sp. nov., orange and *Phakellia pallida* sp. nov., purple.

Within the *P. robusta* species complex we identified three different clades (Clade I-III) with varying degrees of support depending on the genetic marker used (Figs. 9–10). Clade I, contained specimens of *P. robusta* from Røst Reef, Kerry Heads Reef, Korsfjorden, Denmark Strait and the Cantabrian Sea and specimens from Hatton Rockall Bank (HRB) (resequenced in this study), with moderate to strong support for both *cox1* and *28S* (BS = 82 and 99, respectively); Clade I was sister to the group formed by Clade II and Clade III. Clade II, with a weak to strong support value for *cox1* and *28S* (BS = 79 and 96, respectively), was composed by all specimens of *Phakellia pusilla* sp. nov. sequenced in our study, which were collected from the Mediterranean, Cantabrian Sea, and the Canary Islands, which was further subdivided (*cox1* BS = 72), separating Atlantic–Mediterranean specimens from those restricted to the Mediterranean. Clade III, was composed of members of *Phakellia pallida* sp. nov., sequenced in our study, from the Mediterranean and Azores, but also included specimens sequenced by Taboada et. al (2022) collected from the Cantabrian Sea. Clade III showed a moderate support value for both *cox1* and *28S* (90 and 84, respectively).

**Figure 10.**
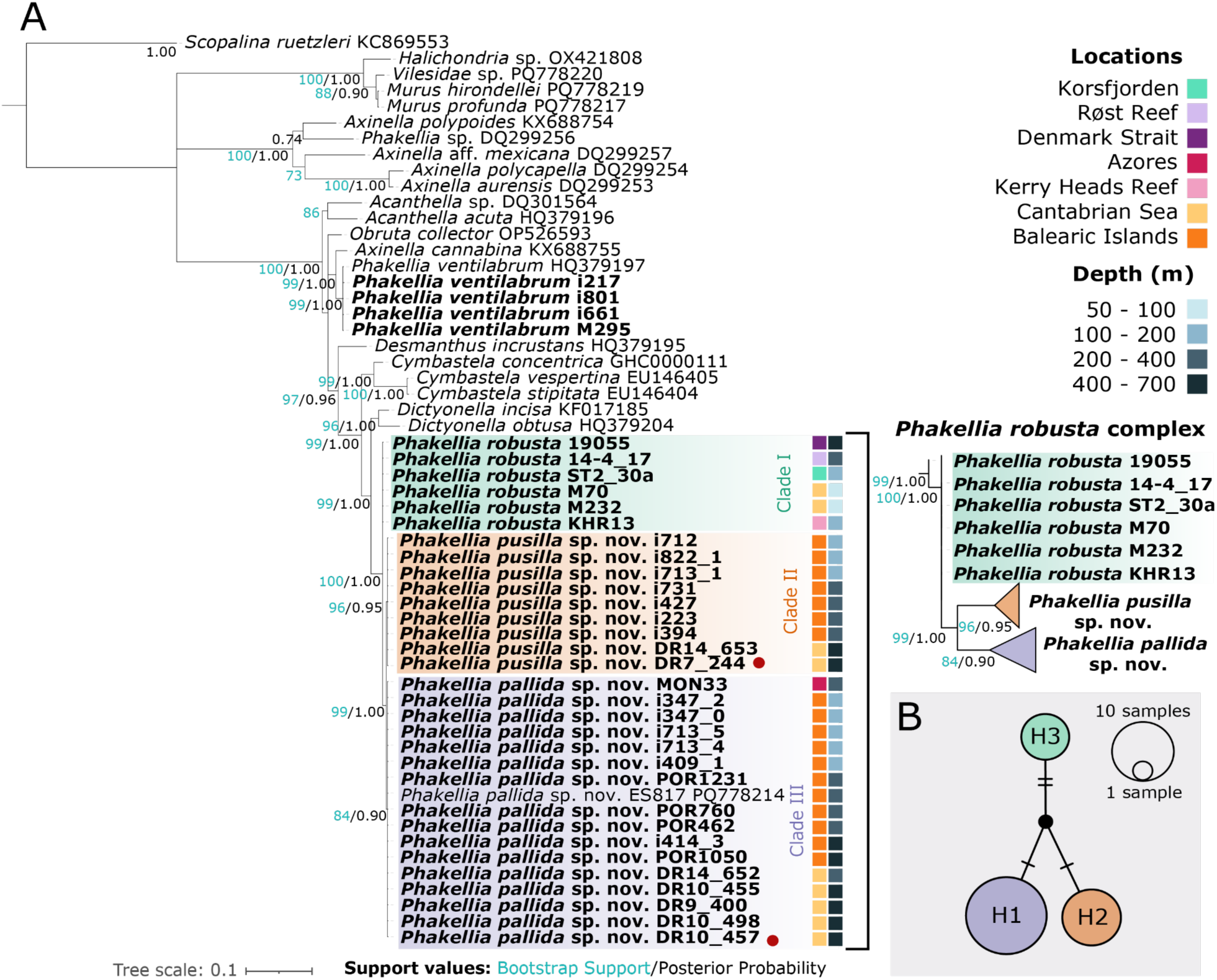
A: Bayesian 50% majority-rule consensus tree inferred with MrBayes based on the *28S* rRNA gene. Numbers at nodes indicate Maximum Likelihood bootstrap support (1,000 replicates; green) followed by Bayesian posterior probabilities (black). Only support values with BS > 70% or PP > 0.70 are shown. Genbank accession numbers appear after each taxon name. Specimens sequenced in our study are in bold. Colored squares show both sampling locations and depth. Red dots mark the individuals designated as type specimens. A magnified view of the phylogenetic tree showing *Phakellia robusta* complex is displayed on the right. B: Haplotype network based on the *28S* for the *Phakellia robusta* complex. Circles are proportional to the number of individuals for each haplotype. Cross lines symbolize the mutational steps and missing inferred haplotypes are in black. Each color represents a different species: *Phakellia robusta* green, *Phakellia pusilla* sp. nov., orange and *Phakellia pallida* sp. nov., purple.

The *cox1* and *28S* (C1–D2) haplotype networks constructed for the *P. robusta* complex identified six and three haplotypes, defined by six and four polymorphic sites, respectively (Figs. 9B, 10B). The number of haplotypes differed among species depending on the marker used, ranging from three (*cox1*; H4-H6) to one (*28S;* H3) in *P. robusta*, and from two (*cox1*; H2-H3) to one (*28S*; H2) in *P. pusilla* sp. nov., while the third lineage was represented by a single haplotype (H1) in both datasets corresponding to *P. pallida* sp. nov.. The two markers revealed contrasting genealogical structures across the complex. In the mitochondrial *cox1* network (Fig. 9B), *P. pallida* sp. nov. (H1) was completely isolated, separated by two mutational steps from an inferred haplotype. In contrast, haplotypes belonging to *P. robusta* (H4–H6) and *P. pusilla* sp. nov. (H2, H3) exhibited a closer relationship, connected with only 1 or 2 mutational steps between them. Conversely, the nuclear *28S* network demonstrated each species defined by a single, unique haplotype (Fig. 10B). These three nuclear haplotypes radiated from a central inferred haplotype, separated by one (*P. pallida* sp. nov. and *P. pusilla* sp. nov.) to two (*P. robusta*) mutational steps.

Interspecific and intraspecific genetic distances were compared for the three species of the *P. robusta* species complex. The three lineages showed interspecific distances in *cox1* between 1.0 and 1.7%, while intraspecific variation observed within each species amounted for 0.27 - 0.48%. For the *28S*, interspecific genetic variation was lower than for the *cox1* (0.2 - 0.5%), also exceeding the maximum intraspecific variation observed within each species, which was 0.0 % for the three species (see Supplementary Table 4).

## Discussion

### Pseudocryptic speciation in Bubarida

In this study we describe two new species of the genus *Phakellia* from the NE Atlantic (Cantabrian Sea and Canary Islands) and the Mediterranean Sea (Balearic Islands), through integrative taxonomy with morphological and molecular approaches. We describe *Phakellia pallida* sp. nov., and *Phakellia pusilla* sp. nov., which are now recognized as new distinct species belonging to the same evolutionary lineage as *Phakellia robusta*. The constructed phylogenies for the *cox1* and *28S* recovered *Phakellia robusta*, *P. pallida* sp. nov. and *P. pusilla* sp. nov. as a closely related species with extremely similar morphological features and small genetic divergences, suggesting that these taxa constitute a species complex (Figs. 9–10). All three species share the general fan-shaped growth form anchored to the substrate by a peduncle, a similar lamellar organization, skeleton patterns in a renieroid organization, and the same monactinal and diactinal spicules with comparable measurements, typical of the genus (Topsent, 1904, 1925) (Figs. 2–7). Such a combination of morphological resemblance and genetic differentiation that is characteristic of species complexes has been previously documented in other sponges, where molecular studies have increasingly helped to uncover cryptic diversity within morphologically conservative taxa (Xavier *et al*., 2010; Swierts *et al*., 2017).

The low levels of genetic divergence observed in both markers and the close phylogenetic relationships among the three taxa are consistent with a recent evolutionary diversification event. Nevertheless, the interspecific variation observed in the *cox1* marker, ranging from 1.0 to 1.7% among the three clades corresponding to *P. robusta*, *P. pusilla* sp. nov. and *P. pallida* sp. nov. (2–5 bp differences), supports their recognition as distinct species within the context of sponges. Small interspecific genetic distances have been previously reported in sponges: for instance, between two cryptic clades of *Suberites diversicolor* (Becking *et al*., 2009), genetic distances of only 0.38% were obtained (Becking *et al*., 2013), although for other cryptic lineages, percentage of nucleotide divergences (*p*-distances) were around 8–10% (e.g. Blanquer & Uriz, 2007; Xavier et al., 2010). Low nucleotide divergences in sponge *cox1* sequences are easily explained given that mitochondrial DNA in early–branching metazoans has lower nucleotide substitution rates due to highly efficient DNA repair mechanisms (Hellberg, 2006; Huang *et al*., 2008; Wang & Lavrov, 2008) (Supplementary Table 4). In contrast, the *28S* marker, despite placing these species in different clades (Fig. 10), showed lower resolution than *cox1*, with lower interspecific variation (0.2 - 0.5%) (2-3 bp differences) (Supplementary Table 4) which is likely due to the recent divergence of these lineages and the limited resolution of slowly evolving markers typically observed in Porifera (Morrow *et al*., 2012).

A detailed examination of both internal and external morphology revealed a set of subtle but consistent characters which, when considered as a whole, facilitate the delineation of the species. Although the three different species presented the same spicular types (including mono- and diactinal spicules), *P. robusta* showed in general wider monactinal and diactinal spicules than *P. pusilla* sp. nov. and *P. pallida* sp. nov. when comparing individuals at similar depths (Table 2; Supplementary Fig. 2). In contrast *P. pallida* sp. nov. and *P. pusilla* sp. nov. showed similar spicule lengths and widths at comparable depths for all spicule types (Table 2). Interestingly, increases in the width of both monactinal and diactinal spicules was positively correlated with depth in most taxa studied, with wider spicules ocurring at greater depths, particularly in Mediterranean populations of *P. pallida* sp. nov. and *P. pusilla* sp. nov., as well as in the Atlantic for *P. robusta* (Supplementary Fig. 2; Supplementary Table 2). This pattern is consistent with previous evidence that spicule dimensions may vary with depth and other environmental conditions (Bavestrello *et al*., 1993; Uriz *et al.,* 2003). Most demosponges depend on dissolved silicate for spicule formation, and high silicate availability enables the production of larger spicules, both in length and width (Yourassowsky & Rasmont, 1983; Maldonado et al., 1999: Mercurio et al., 2000). Because dissolved silicate concentrations generally increase with depth (John *et al*., 2022), the increase in spicule width observed here may therefore be partly related to greater silica availability in deeper water layers (Supplementary Fig. 2). However, other environmental factors associated with depth cannot be excluded.

Apart from differences observed at the spicule level, other differences were found when examining in detail the external morphology of the three species studied here. Both *P. pallida* sp. nov. and *P. pusilla* sp. nov. possess markedly thinner lamellae and a more delicate body than *P. robusta*. In addition, *P. pusilla* sp. nov. is characterized by the presence of cribriporal pores, whereas *P. robusta* and *P. pallida* sp. nov. exhibit uniporal pores. The most conspicuous feature distinguishing *P. pallida* sp. nov. is the presence of well-developed macromorphological veins or striations, which are weakly expressed or absent in the other two species. Moreover, while *P. robusta* has never been reported to develop a cup-shaped morphotype (Topsent, 1904, 1928; Ferrer Hernández, 1914; Santín *et al*., 2018), both newly described species can adopt either fan-shaped or cup-shaped forms, a shared phenotypic plasticity that may lead to confusion with *P. ventilabrum* or *P. robusta* during field identification (Díaz *et al*., 2021). In fact, recent molecular phylogenetic analyses have shown that several specimens historically assigned to *Phakellia* were misidentified. These include material from the Topsent collection from North Atlantic localities (Topsent, 1890, 1892; Boury-Esnault *et al*., 1994), as well as specimens from the Cantabrian Sea studied more recently (Taboada *et al*., 2022). Díaz et al. (2025) clarified that the specimens identified as *Phakellia hirondellei* in Taboada et al. (2022) from the Cantabrian instead represent a different species nominated as *Phakellia* aff. *robusta*. These specimens, which cluster within the *P. robusta* clade in our phylogenetic analysis, correspond in fact to the new species described here as *Phakellia pallida* sp. nov. In addition, some individuals also studied in Taboada at al. (2022) were identified as potential *P. robusta* hybrids from the Cantabrian Sea (DR7-244 (accession number MT506138), DR14-651 (accession number MT506139) and DR14-653 (accession number MT506140)) indeed constitute the second species described in our study, *Phakellia pusilla* sp. nov., although the nature of the other hybrids identified in the study cannot be discarded. Taken together, these subtle but consistent morphological differences, combined with their genetic differentiation, support the interpretation of these taxa as pseudocryptic rather than truly cryptic species.

### Ecology and biogeography of newly described Phakellia species

The two newly described *Phakellia* species showed overlapping distributions, both occurring across the Atlantic–Mediterranean region. Both species also share bathymetric distribution, spanning from the lower circalittoral down into the upper bathyal (upper slope) (132 to 674 m) although we found *P. pusilla* sp. nov. more frequently at shallower circalittoral depths than *P. pallida* sp. nov (Table 1). This substantial overlap in habitat conditions may help explain their overall morphological resemblance, as similar environmental conditions may have constrained morphological divergence (Blanquer & Uriz, 2007). Consequently, their phenotypic resemblance could reflect that they speciated recently. It is interesting to note that in the Cantabrian Sea, specimens of both species were generally collected between 340 and 660 m depth, whereas most Mediterranean specimens occurred at shallower depths (140–400 m) with a single individual collected below 600 m. This pattern could reflect depth- and circulation-related differences in dispersal, as well as regional differences in productivity and food sources. In the Mediterranean Sea, and particularly in the highly oligotrophic waters around the Balearic Islands, benthic communities may depend strongly on energy ultimately derived from primary production in the euphotic zone, with surface-derived energy extending its influence into mesophotic habitats (Zabala & Ballesteros, 1989; Díaz *et al*., 2024). By contrast, the Cantabrian Sea margin receives substantial river-sourced particulate inputs, and canyon-mediated processes facilitate the transport of this material towards deeper habitats (Rumín-Caparrós *et al*., 2016). This enhanced supply of material may contribute to the apparently deeper distribution of the species in this region. However, this pattern could also reflect sampling bias, and further sampling in these areas is needed to determine whether the observed depth differences are robust.

Our results suggest that *P. robusta* is likely a boreal species, with its southernmost confirmed records in the Cantabrian Sea, and is probably absent from the Mediterranean Sea. Originally, *P. robusta* was described from the Shetland Islands, located in the Scotland Archipelago (Bowerbank, 1866). Since then, this species has been cited in the same area (Stephens, 1921; Alander, 1942) and in other parts of the Atlantic Ocean, including the Cantabrian Sea (Ferrer Hernández, 1914; Ríos *et al*., 2017; Sánchez *et al*., 2017; Taboada *et al*., 2022), Azores (Topsent, 1904, 1928), Madeira (Topsent, 1928), Canary Islands (Cruz, 2002) and in the Mediterranean Sea (Topsent, 1925; Vacelet, 1969; Maldonado, 1992; Santín *et al*., 2018) at depths ranging from 20 m to 1,229 m. In our study, we report the first record of *P. robusta* in Iceland, based on an individual collected from the Denmark Strait at approximately 500 m depth (191055). This specimen clustered with other *P. robusta* samples, differing by a single nucleotide in the *cox1*, which may reflect population-level differences (Fig. 9). The records of *P. robusta* found throughout the Mediterranean Sea may actually correspond to individuals of *P. pallida* sp. nov. or *P. pusilla* sp. nov., as it happened for the individuals, we studied from the Balearic Islands: i347_2, i405, i409, i417, i712, i731 and POR760 (Figs. 9–10), which were previously misidentified as *P. robusta* (Díaz *et al.,* 2021). The specimen MON33 from Josephine Bank, off the coast of the Iberian Peninsula (NE Atlantic), originally described by Topsent (1928) as *P. robusta* var. *hirondellei* and later reassigned to *P. robusta* by Díaz *et al.,* (2025), was re-examined in this study. Both morphological observations and molecular analyses (*cox1* and *28S*) confirm that this specimen should be assigned to *P. pallida* sp. nov. (Figs. 9–10; Table 2). Some southern records currently attributed to *P. robusta* (e.g., from the Canary Islands; Cruz 2002) may instead correspond to *P. pusilla* sp. nov., which is also present in the region (Table 1) These reassignments suggest that the southernmost distribution of *P. robusta* may be the Cantabrian Sea. However, additional sampling and genetic analyses are needed to confirm these patterns and to determine whether *P. robusta* truly occurs in the Mediterranean and further south of the Cantabrian Sea.

### Phylogenetic hypotheses and intron presence/absence

Similar to previous recent studies including members of the genus *Phakellia* (Taboada et al., 2022; Díaz et al., 2025), our phylogenetic results recovered the genus *Phakellia* as polyphyletic, with some members of *Phakellia* (including *P. robusta, P. pallida* sp. nov., and *P. pusilla* sp. nov.) forming a well-supported clade for *cox1* and appearing as sister clade of another one including an identified member of the genus *Phakellia* and taxa from the genera *Dictyonella* and *Cymbastela*. Importantly, this clade that included members of the genera *Phakellia*, *Dictyonella* and *Cymbastela*, was separated from that including *P. ventilabrum*, the type species of the genus (Bowerbank, 1862), *P. carduus* and *Phakellia rugosa* (Bowerbank, 1866) (Figs. 9–10), as previously found (Taboada *et al*., 2022). Although our results indicate that the clade including *P. robusta, P. pallida* sp. nov., and *P. pusilla* sp. nov. should be considered as an independent evolutionary lineage potentially constituting a new genus, we prefer not to take any taxonomical action to this respect given the limited availability of molecular and morphological data for other *Phakellia* species.

The topological discordance observed between the mitochondrial *cox1* and nuclear *28S* phylogenies highlights distinct sister relationships within the *Phakellia robusta* complex (Figs. 9–10). While *cox1* places *P. robusta* and *P. pusilla* sp. nov. closer with *P. pallida* sp. nov. as the divergent lineage, the nuclear *28S* topology groups *P. pusilla* sp. nov. and *P. pallida* sp. nov. closer together, isolating *P. robusta*. Such cyto-nuclear discordance typically arises from either hybridization or from incomplete lineage sorting during a rapid radiation of the complex (Toews & Brelsford, 2012). Distinguishing between these two evolutionary scenarios will require genome-wide multilocus data, as single-locus markers have lower resolution in resolving these evolutionary processes (Som, 2014; Edelman *et al*., 2019).

Previous studies have described the presence of a group I intron in the *cox1* gene of *P. robusta* (Cranston *et al*., 2021). In the present study, we detected the same type of intron inserted at the same position, although with a shorter length, in some individuals of *P. pusilla* sp. nov. (i417, i629_1_1, i712, i713_3, i822_1) from the Balearic Islands (Fig. 8). In contrast, the intron was not detected in any individual of *P. pallida*. The occurrence of a group I intron in *P. robusta* and in some individuals of *P. pusilla* sp. nov. suggests that the intron was likely acquired by their most recent common ancestor. This hypothesis is further supported by the mitochondrial phylogeny, in which *P. robusta* and *P. pusilla* sp. nov. form a supported clade, whereas *P. pallida* falls outside this lineage (Fig. 8). The absence of the intron in all examined *P. pallida* individuals is therefore consistent with the intron having originated after the divergence of *P. pallida*. The fact that the intron occupies the same position in both species (Fig. 8A) is also consistent with the tendency of sponge group I introns to insert into highly conserved regions (Schuster *et al*., 2017). Given that *P. robusta* and *P. pusilla* are closely related sister species, their *cox1* sequences are highly similar, particularly in conserved regions, which would provide an equivalent target site for intron insertion in both lineages. However, the absence of the intron in some *P. pusilla* sp. nov. individuals and the distinct size of the intron relative to that observed in *P. robusta* complicates this scenario (Fig. 8B). It may suggest either a secondary loss of the intron within *P. pusilla* sp. nov. or alternatively that the common ancestor lacked the intron and that the shared insertion site reflects independent insertion events of two different introns in the two sister species. The latter hypothesis could be explained by a relatively recent acquisition of the group I intron in *P. pusilla* sp. nov., with insufficient time for it to become fixed within the species, potentially due to genetic drift and small effective population sizes as it has been proved to happen in populations of the genus *Daphnia* with absence/presence of introns in different populations (Omilian *et al*., 2008). Although the intron was found only in individuals from Balearic Islands, we also identified other samples from this region without it. A third possibility is hybridization. As suggested by Taboada et al. (2022), the specimens identified here as *P. pusilla* sp. nov. may represent hybrids involving *P. robusta*. Under this scenario, individuals inheriting the maternal *cox1* haplotype from *P. robusta* could also have inherited the intron, explaining its presence in these specimens and its absence in *P. pallida* sp. nov. This study includes a limited number of samples from the Cantabrian Sea, so additional sampling would be necessary to verify whether this intron is indeed absent in individuals from that region as well. However, future research in the mitogenome of *P. pusilla* sp. nov. is needed to elucidate the evolutionary process underlying this polymorphism in the *cox1*.

## Conclusion

This study highlights the importance of recognizing pseudocryptic diversity in deep-sea sponges. The discovery of *Phakellia pallida* sp. nov. and *P. pusilla* sp. nov. within a group historically identified as *P. robusta* demonstrates that species diversity can remain underestimated when species identification relies primarily on morphology. Identifying and resolving pseudocryptic species is essential not only for an accurate assessment of biodiversity, but also for understanding the biogeographical and ecological characteristics of individual species, as the actual functioning and specificities of each species can be masked (Xavier et al., 2010). The distribution of *Phakellia robusta* should therefore be treated with caution, particularly in the Mediterranean Sea, where many records remain to be confirmed using molecular data, as they could in fact correspond to the new species described here. Continued sampling and molecular characterization of both newly collected and historical specimens will consequently be essential to clarify the true distribution and diversity of the *P. robusta* species complex. Finally, genome-wide approaches will be essential to unravel the evolutionary mechanisms driving diversification in this complex.

## Acknowledgments

We thank the crews and scientists aboard the R/V *Ángeles Alvariño*, R/V *Ramón Margalef* and R/V *Miguel Oliver* for their work during the research surveys SponGES0617, INTEMARES_A22B_1019, INTEMARES_A22B_0720, CIRCAROCK, CIRCA-LEBA-0523 and MEDITS, with special thanks to Francesc Ordines, chief scientist of the MEDITS surveys, for his support and coordination during these campaigns.

We are grateful to Antoni Busquets (UIB) for his assistance with SEM analyses. Many thanks to Paco Cárdenas for hosting C.G.S. at the Museum of Evolution, for his valuable comments on the manuscript, and for his help in disentangling the taxonomic history of *Phakellia robusta*. We are also indebted to Christine Morrow for providing valuable samples for this study.

## Supplementary Data

**Supplementary Figure 1:**
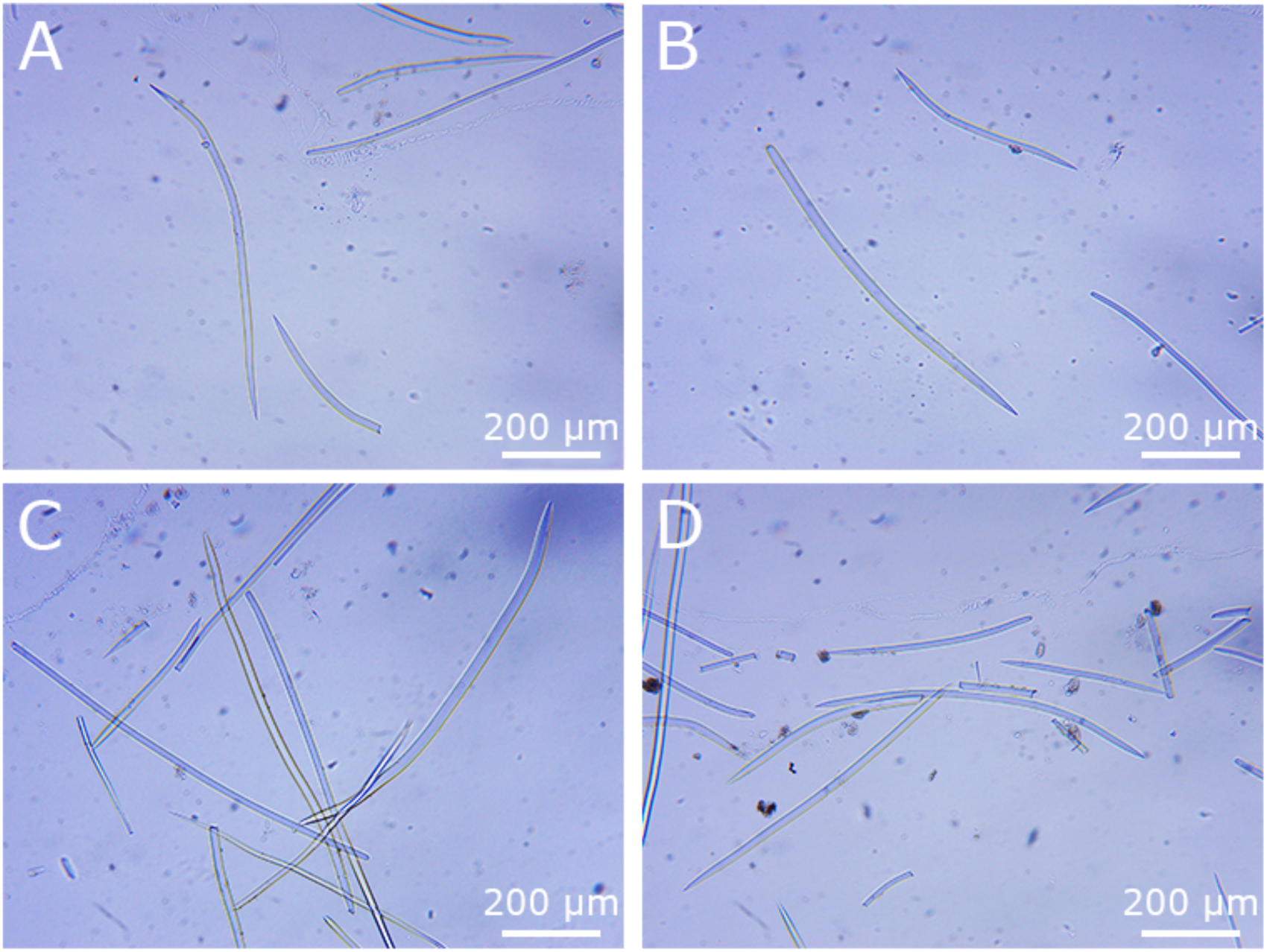
Optical microscope images of *Phakellia robusta* holotype 420. A. Diactinal (Oxea) and monactinal (stylota) spicules. B. Monactinal (Stylota) and monactinal (Oxeota). C. Spicule mixture. D. Spicule mixture.

**Supplementary Figure 2:**
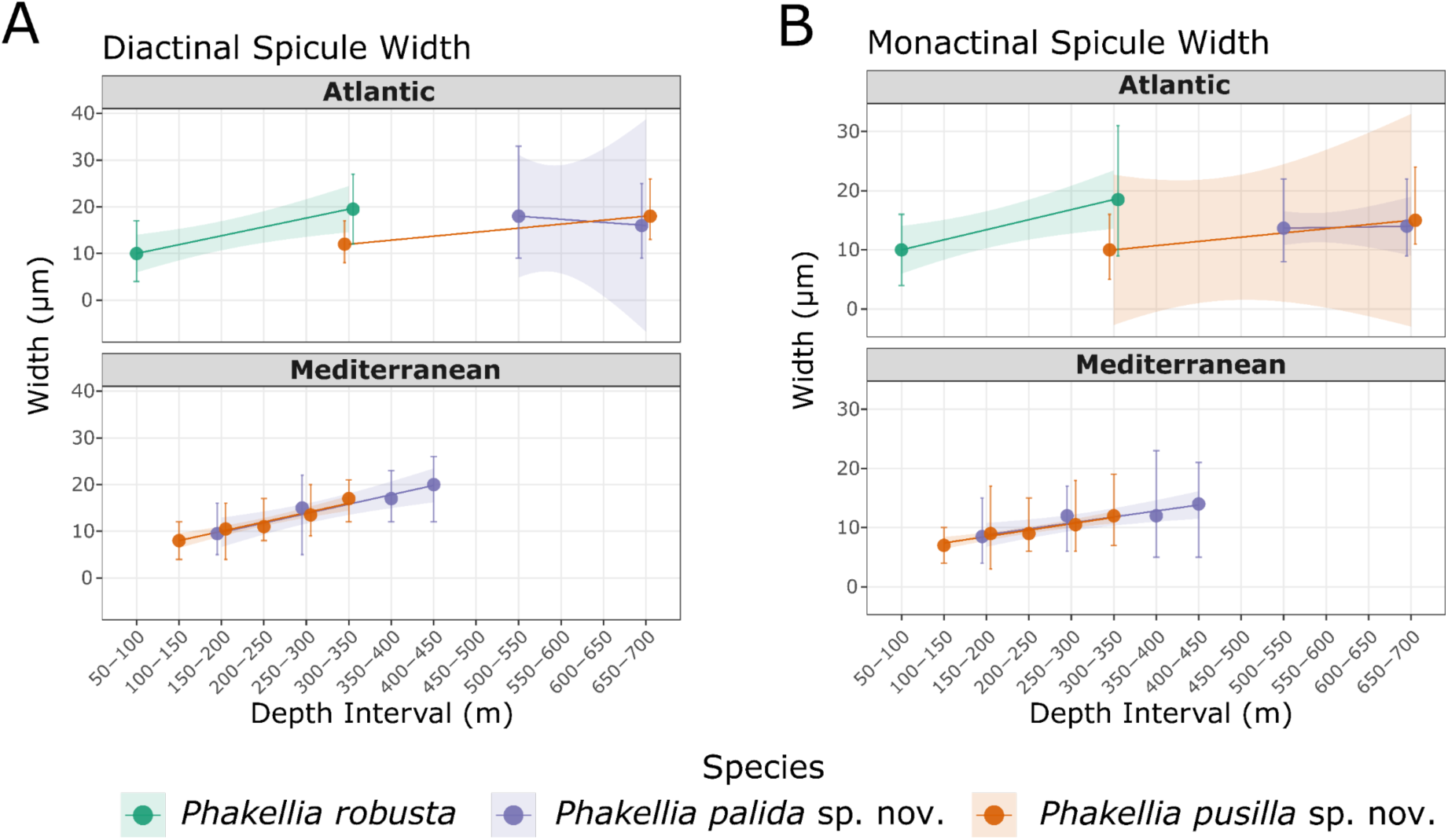
*Phakellia* spp. spicule width variation. Morphometric variation in the spicules width of *Phakellia robusta* (green), *Phakellia pallida* sp. nov. (purple) and *Phakellia pusilla* sp. nov. (orange) positively related to depth. The points represent the consolidated mean value of the specimens grouped within each bathymetric interval, while the vertical error bars indicate the absolute minimum and maximum morphometric ranges observed. The coloured solid lines represent independent linear regression models (y ∼ x) calculated for each species, with shaded bands indicating the corresponding 95% confidence intervals. A. Diactinal spicules width variation across standardised depth intervals of 50 m shown for Atlantic and Mediterranean individuals. B. Monactinal spicules width variation across standardised depth intervals of 50 m shown for Atlantic and Mediterranean individuals.

**Supplementary Figure 3:**
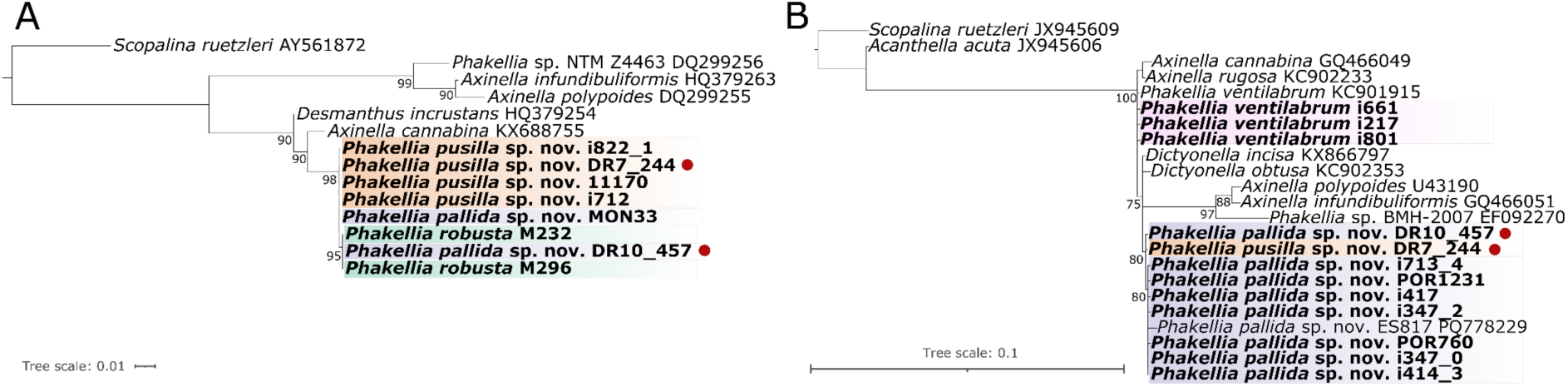
Phylogenetic tree constructed with RAxML based on the *28S* (D3-D5 region) (A) and *18S* (First fragment) (B) using Maximum Likelihood (ML). At each node ML bootstrap supports (1000 bootstrap replicates) are shown (only bootstrap values > 70 are reported in the tree). Genbank accession numbers appear after each taxon name. Specimens sequenced in our study are in bold. Red dots mark the individuals designated as type specimens.

**Supplementary Figure 4:**
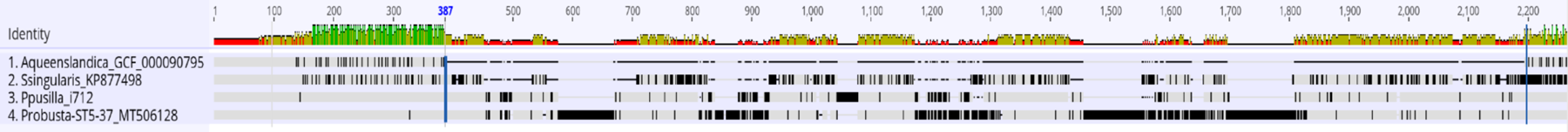
Intron alignment. Alignment done with *cox1* sequences from *Amphimedon queenslandica*, *Stupenda singularis*, *Phakellia robusta* and individual i712 from *Phakellia pusilla* sp. nov. The insertion site is indicated in bold at position 387 bp, extending to approximately 2,200 bp.

- **Supplementary File 1: *cox1* alignment.** 440 bp and 85 sequences. Best-fit model HKY + I + G.

- **Supplementary File 2: *28S* (C1-D2) alignment.** 828 bp and 57 sequences. Best-fit model GTR + I + G.

- **Supplementary File 3: *28S* (D3-D5) alignment.** 569 bp and 14 sequences. Best-fit model GTR + I.

- **Supplementary File 4: *18S* (1° fragment) alignment.** 537 bp and 23 sequences. Best-fit model K80 + I.

- **Supplementary Table 1: Metadata.** Metadata of all the sponges used in the study, including codes of the sequences uploaded in NCBI in sheet 2.

- **Supplementary Table 2: Statistical parameters of spicule analysis.** Summary of linear regression analyses evaluating the relationship between spicule width (µm) and continuous collection depth (m) across *Phakellia* species and geographic basins. *n*, number of analyzed specimens; *β*, regression slope (Estimate); SE, standard error of the slope; *R²*, coefficient of determination; *p*, significance level of the slope parameter (\*\*\**p* < 0.001, *p* < 0.01, \**p* < 0.05, *ns*: not significant, *p* ≥ 0.05).

- **Supplementary Table 3: Spicule measurements.** Details table of spicule measurements for all individuals included in Table 2.

- **Supplementary Table 4: Interspecific genetic distances.** Interspecific and intraspecific mean pairwise genetic distances calculated for the species used in the phylogeny using the *cox1* and *28S* alignments. Each sheet corresponds to a different gene respectively. Values represent the evolutionary distances computed under the Kimura 2-parameter (K2P) model in the interspecific distances whereas intraspecific distances are reported as maximum and mean values.

## Data Availability Statement

The DNA sequence data generated in this study have been deposited in GenBank and are available under the accession numbers QB057393-QB057395; QB057355-QB057392; QB044867-QB044901; QB044902-QB044910; QB053226-QB053237. The sequence alignments used for phylogenetic analyses, specimen metadata, spicule measurements, statistical analyses and genetic distance data are provided in the Supplementary Material. All other data supporting the findings of this study are available within the article and its Supplementary Material.

