## Supplementary figures and images for "Pseudocryptic diversity within the *Phakellia robusta* species complex: description of *Phakellia pallida* sp. nov. and *Phakellia pusilla* sp. nov"

### Supplementary Figure 1

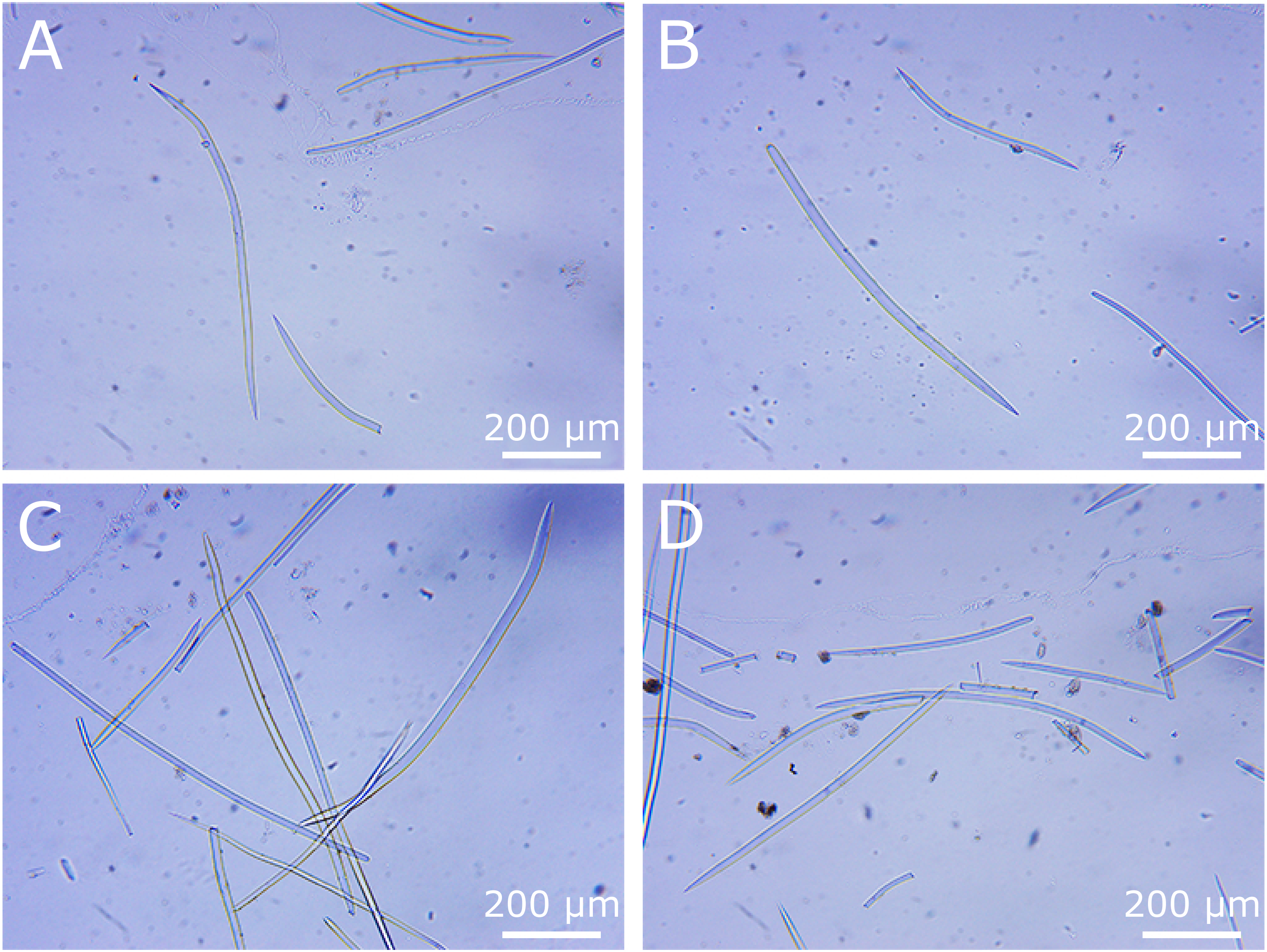

### Supplementary Figure 2

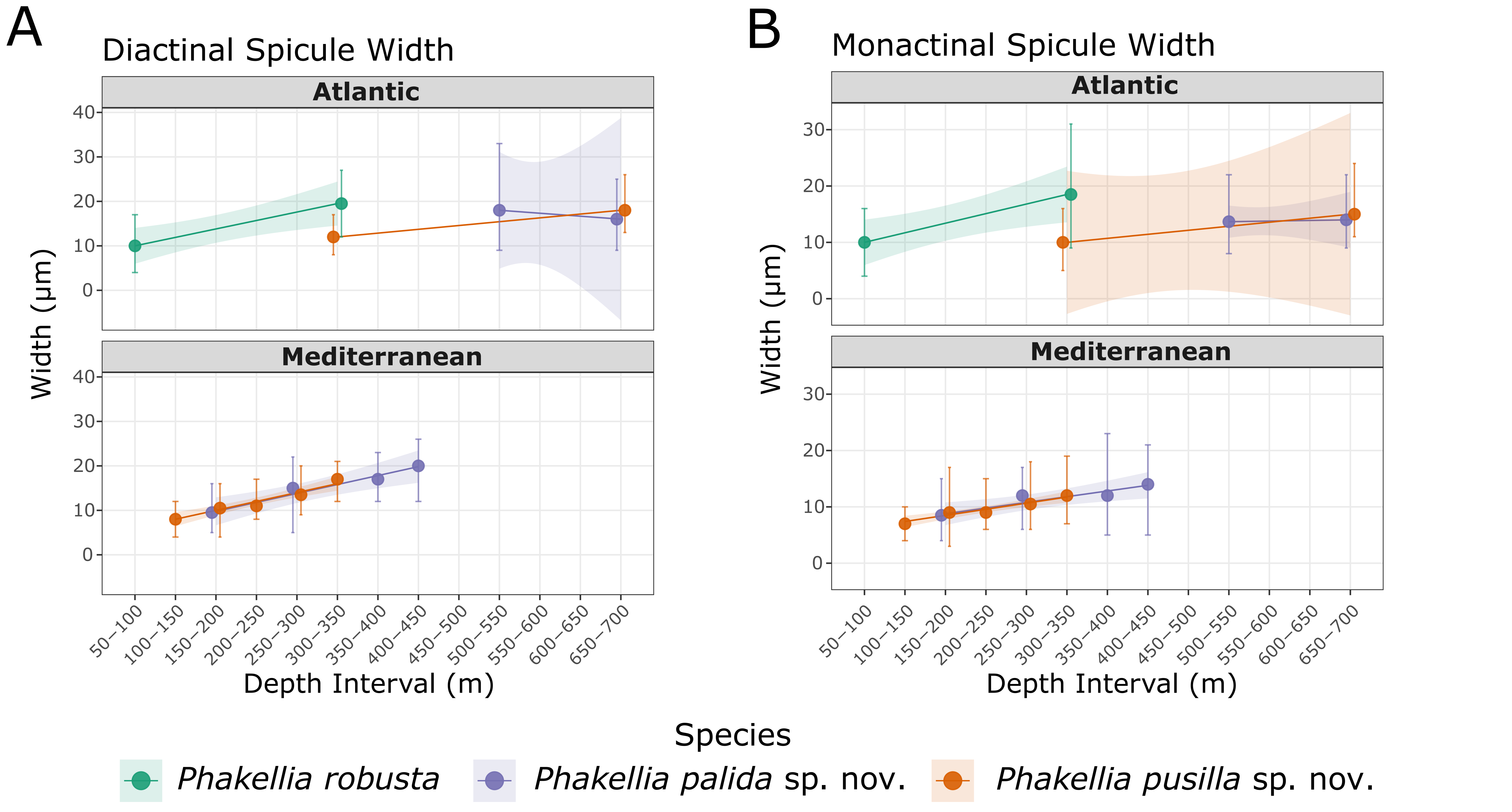

### Supplementary Figure 3

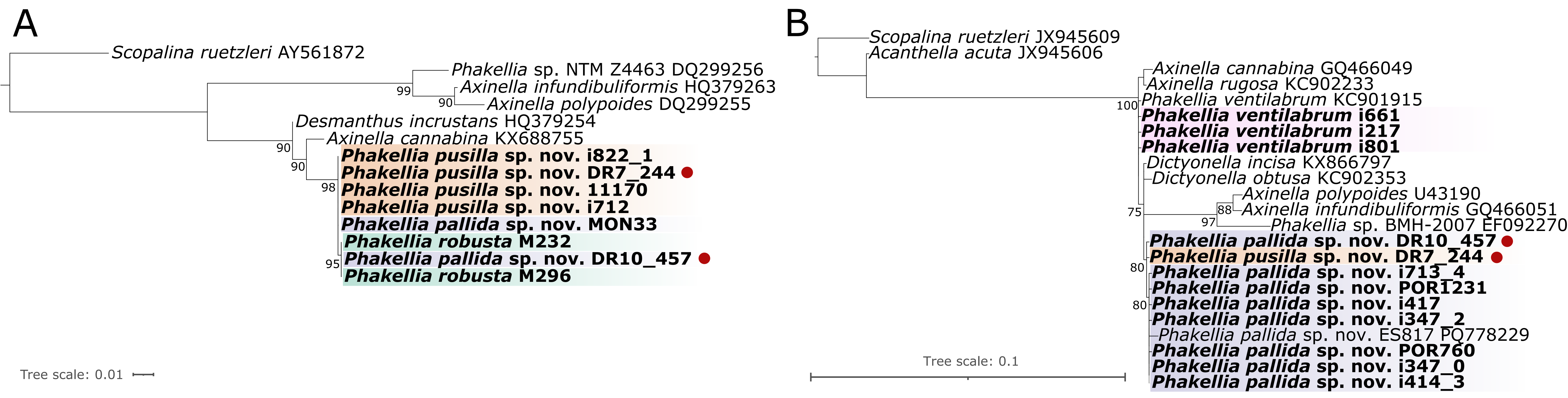

### Supplementary Figure 4

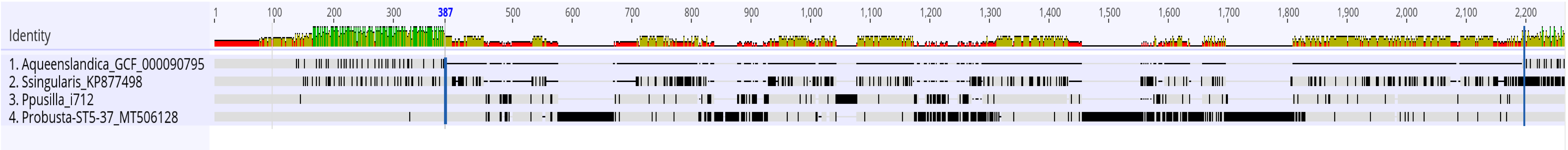
